# TICToK: A comprehensive knowledgebase of tattoo ink chemicals and investigation of their associated toxicities and regulations

**DOI:** 10.1101/2025.08.02.668261

**Authors:** Nikhil Chivukula, Shreyes Rajan Madgaonkar, Shambanagouda Rudragouda Marigoudar, Krishna Venkatarama Sharma, Vimal Kishore, Ajay Vikram Singh, Areejit Samal

## Abstract

Tattoos and permanent make-up, involving dermal injection of chemicals, are increasingly popular, with 30-40% of young adults in North America and Europe having at least one tattoo. Recent studies have suggested a possible association between tattooing and health risks, including skin cancers and lymphomas, although no definitive link has yet been established. This study comprehensively catalogs tattoo ink chemicals and investigates their potential adverse effects, addressing the urgent need for greater understanding of tattoo-related health concerns. First, 364 unique tattoo chemicals were identified from various scientific and regulatory sources, with nearly half functioning as pigments. Hazardous chemicals were identified, revealing carcinogens, endocrine disruptors, neurotoxicants, and dermal toxicants. A regulatory analysis based on key EU regulations, including harmonised classifications under Classification, Labelling and Packaging (CLP) regulation, Restriction Entry 75 of the REACH regulation, Cosmetic Products Regulation (CPR), and SVHC candidate list, revealed existing regulatory coverage of tattoo ink chemicals. Curated chemical-disease associations highlighted that some tattoo chemicals are known to cause dermatitis. Further, diverse toxicological information, including experimental results from REACH dossiers, were integrated to construct stressor-AOP network linking 151 chemicals to 362 AOPs, revealing potential carcinogenic mechanisms associated with tattoo ink chemicals. A systems biology approach revealed potential immunomodulatory effects associated with these chemicals. Finally, all findings have been made available through the online database **T**attoo **I**nk **C**hemicals and associated **To**xicities **K**nowledgebase (TICToK; https://cb.imsc.res.in/tictok), which can support risk assessment and sustainable tattoo practices.

**Graphical Abstract:** 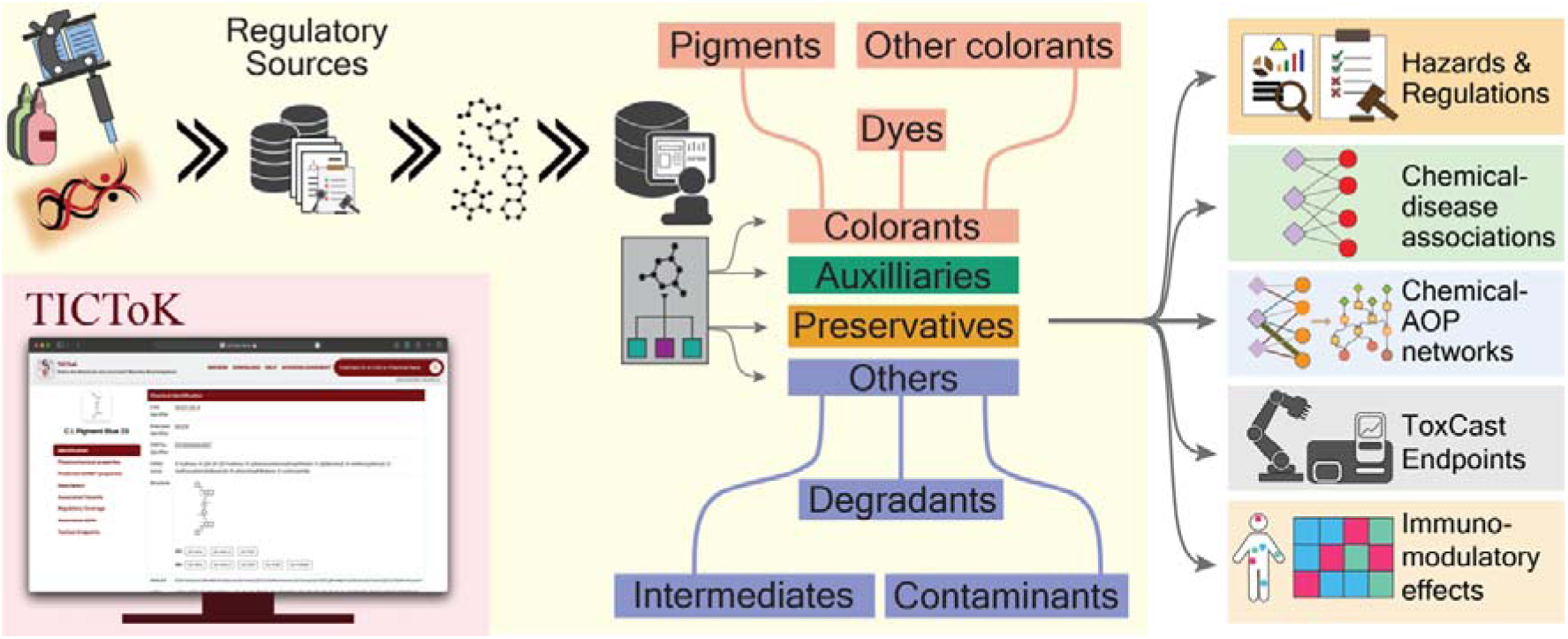

## 1. Introduction

Tattooing is the practice of depositing pigment into the dermal layer of the skin to create permanent body art (Sizer, 2020). Permanent make-up (PMU) is a closely related technique involving micropigmentation of facial features to provide long-lasting cosmetic effects (Ghafari et al., 2024). Recent surveys indicate that nearly 30-40% of young adults in Europe and North America have at least one tattoo (Laumann and Derick, 2006; Kluger et al., 2019; Ghafari et al., 2024; IARC, 2025). Beyond these specific regions, a recent review reports varying rates of tattoo prevalence worldwide (Palese and Valent, 2025). Thus, it is imperative to gain a systematic understanding of the chemicals present in tattoo inks and their associated adverse effects, to aid in risk assessment and support sustainable and safe tattoo practices.

Modern tattoo inks incorporate a wide range of chemicals to improve color vibrancy, stability, and longevity (Laux et al., 2016; Randhawa, 2024). The European Commission’s Joint Research Centre (EU JRC) published a series of reports titled ‘Safety of tattoos and permanent make-up’, systematically cataloging the chemicals present in tattoos, through interactions with various stakeholders, as well as a comprehensive literature survey (Piccinini et al., 2015, 2016a, 2016b). The report further categorized the diverse chemicals into main classes such as colorants, auxiliaries, and preservatives to denote their functional role in tattoo inks (Piccinini et al., 2015). Similarly, the Australian National Industrial Chemicals Notification and Assessment Scheme (NICNAS) published comprehensive reports analyzing the tattoo and PMU inks marketed within Australia (National Industrial Chemicals Notification and Assessment Scheme (NICNAS) and Australian Government, Department of Health, 2018, 2017). Additionally, the NORMAN Suspect List Exchange (NORMAN-SLE) (Mohammed Taha et al., 2022) published a curated list of suspect chemicals in tattoo inks to support their monitoring and regulatory surveillance, sourcing its contents from the regulated ingredients list for tattoo ink and PMU established under Appendix 13 of Commission Regulation (EU) 2020/2081, which amended Annex XVII of REACH regulation (https://eur-lex.europa.eu/eli/reg/2020/2081/oj/eng). Each of these initiatives independently covered a limited subset of the chemicals occurring in tattoo inks and revealed little consensus on the full spectrum of compounds currently present in products across the globe.

The JRC reports further compiled the different hazards associated with these tattoo chemicals (Piccinini et al., 2016a, 2016b). Notably, the report highlighted the lack of risk assessment that takes into account their injection into human body for long term permanence, as they were originally produced for other applications, such as paints (Piccinini et al., 2016b). Growing evidence of adverse health effects associated with tattoo chemicals has spurred further research efforts (Giulbudagian et al., 2024), and different studies have raised concerns of elevated risks of dermatological conditions and lymphomas associated with tattoos (Bassi et al., 2014; Paprottka et al., 2014; Nielsen et al., 2024; Rigali et al., 2024; Brusasco et al., 2025; Clemmensen et al., 2025; McCarty et al., 2025). Despite growing evidence of hazards associated with tattoo chemicals, the aforementioned reports revealed gaps in their regulation. To address the growing safety concerns, the German Federal Institute for Risk Assessment (BfR) has proposed a set of minimal toxicological requirements to evaluate the risks posed by tattoo ink chemicals (BfR, 2021). These include assessments for eye and skin irritation, skin sensitization, phototoxicity, and genotoxicity (BfR, 2021). In line with this report, Pradeep *et al*. (Pradeep et al., 2025) curated a list of tattoo pigments and explored the genotoxic potential of these chemicals by utilizing experimental evidence from the REACH registration data and various QSAR models, demonstrating the utility of new approach methodologies (NAMs) in tattoo ink risk assessment. Moreover, the study highlighted the necessity of curating chemical information and associated hazards from diverse sources to enable gap-filling for data-poor chemicals (Pradeep et al., 2025). Such efforts not only aid in developing better hazard identification strategies but also support the identification of safer alternatives to known toxic compounds, thereby promoting safer tattoo practices.

Consequently, in this study, multiple regulatory and scientific resources were first leveraged to systematically compile and curate tattoo ink chemicals and annotate their respective functions. Subsequently, associated hazards were identified by consolidating information on known carcinogens, endocrine disruptors, neurotoxicants, and chemicals known to cause skin or eye irritation, skin sensitization, and genotoxicity. The regulatory coverage of these chemicals was then evaluated with respect to EU REACH Restriction Entry 75 and the EU cosmetic products regulation. To further understand their biological relevance, disease associations were extracted from the Comparative Toxicogenomics Database (CTD). Moreover, diverse toxicological endpoints, including those from REACH chemical dossiers, were integrated using the Adverse Outcome Pathway (AOP) framework, resulting in a stressor-AOP network that highlights potential toxicity mechanisms underlying tattoo chemical-induced toxicities. Further, a systems biology-based analysis was undertaken to examine the potential immunomodulatory effects of these tattoo chemicals in the skin. Finally, all results and curated data have been compiled into an interactive online resource, the Tattoo Ink Chemicals and their associated Toxicities Knowledgebase (TICToK), accessible at https://cb.imsc.res.in/tictok/, to support future research and regulatory efforts in this domain.

## 2. Material and Methods

### 2.1. Compilation and curation of tattoo ink chemicals

In this study, a comprehensive list of chemicals in tattoo inks was systematically compiled from the following resources: (i) NORMAN Suspect List Exchange (NORMAN-SLE) (Mohammed Taha et al., 2022) (https://www.norman-network.com/?q=suspect-list-exchange) that provides a curated list of suspect chemicals in tattoo inks (last accessed on 27 May 2025); (ii) CompTox Chemical Dashboard’s (Williams et al., 2017) chemical list on tattoo inks (https://comptox.epa.gov/dashboard/chemical-lists/TATTOOINK) (last accessed on 27 May 2025); (iii) the United States Chemical and Products Database (CPDat) (Dionisio et al., 2018) list of chemicals reported to be present in tattoo inks marketed in the US, which is now hosted via the United States Environment Protections Agency’s ChemExpo knowledgebase (https://comptox.epa.gov/chemexpo/get_data/) (last accessed on 27 May 2025); (iv) the European Union Joint Research Commission (EU JRC) report titled ‘Safety of tattoos and permanent make-up’ (Piccinini et al., 2015); (v) the reports by Australian Industrial Chemicals Introduction Scheme (formerly known as the National Industrial Chemicals Notification and Assessment Scheme (NICNAS)) titled ‘Characterisation of tattoo inks used in Australia’ (National Industrial Chemicals Notification and Assessment Scheme (NICNAS) and Australian Government, Department of Health, 2018) and ‘Investigation of the composition and use of permanent make-up (PMU) inks in Australia’ (National Industrial Chemicals Notification and Assessment Scheme (NICNAS) and Australian Government, Department of Health, 2017); (vi) the internally compiled lists of banned and non-banned pigments in the European Union (EU) (Supplementary Information). The detailed descriptions of each chemical list are provided in the Supplementary Information. Notably, the CompTox chemical list on tattoo inks and the NORMAN-SLE entry for tattoo inks both derive from the same primary source, Appendix 13 to Restriction Entry 75 of Annex XVII to the REACH Regulation as amended by the Commission Regulation (EU) 2020/2081 (European Parliament and European Council, 2020), representing an identical dataset rather than independent compilations. All chemicals reported across these resources were included in the compiled list without any exclusions.

Thereafter, chemical information was standardized by mapping them to corresponding Chemical Abstracts Service Registry Numbers (CASRN) (https://commonchemistry.cas.org/), PubChem chemical identifiers (https://pubchem.ncbi.nlm.nih.gov/), and Distributed Structure-Searchable Toxicity (DSSTox) (Grulke et al., 2019) identifiers (DTXSID) by the US Environmental Protection Agency (USEPA), resulting in the identification of 364 unique chemicals (Table S1). Furthermore, the chemical structures corresponding to these tattoo chemicals were obtained from PubChem (Supplementary Information). In particular, only 322 out of the 364 chemicals had structural information available in PubChem (Table S1). For these chemicals, their corresponding chemical classifications, such as Kingdom, Superclass, and Class, were obtained using ClassyFire (Djoumbou Feunang et al., 2016) (http://classyfire.wishartlab.com/). For chemicals lacking such classification, structural definitions were used to manually assign them to appropriate Kingdoms (Djoumbou Feunang et al., 2016). Through this process, 276 chemicals were categorized as organic compounds and 46 as inorganic compounds (Table S1). Among the remaining chemicals, 19 were identified as mixtures, 15 as polymers, and 6 lacked a defined chemical structure (Supplementary Information; Table S1).

### 2.2. Classification of tattoo ink chemicals based on their reported functions

Tattoo inks are complex formulations of chemicals, comprising pigments suspended in binders and solvents, and additives like preservatives to protect against bacterial contamination (Dirks, 2015; Giulbudagian et al., 2020; Piccinini et al., 2015). The reports by JRC (Piccinini et al., 2015) and Australian NICNAS (National Industrial Chemicals Notification and Assessment Scheme (NICNAS) and Australian Government, Department of Health, 2018, 2017) classify the tattoo ink chemicals into broad categories such as colorants (pigments, dyes), auxiliaries (binders, solvents, etc.), and preservatives, based on their reported function in tattoo inks. Further, CPDat provides functions of these chemicals in the tattoo inks, and the lists of banned and non-banned pigments in the EU provide a compiled list of pigments. In this study, the compiled list of 364 tattoo ink chemicals were therefore categorized into ‘colorants’, ‘auxiliaries’ and ‘preservatives’ based on their reported function(s) from each of these sources (Table S2). For chemicals lacking such information, the CompTox Dashboard’s chemical lists titled ‘AZODYES’ and ‘CIDYES’, were relied upon to identify the colorants, and ‘PARISIII’ and ‘WIKISOLVENTS’ were relied upon to identify the auxiliaries (Table S2). In addition, chemical lists from books titled ‘Industrial Organic Pigments’ (Hunger and Schmidt, 2018) and ‘Industrial Dyes’ (Hunger, 2002) were relied upon to identify colorants among the 364 chemicals (Supplementary Information; Table S2).

In tattoo inks, pigments are the preferred colorants as they are insoluble in the medium they are incorporated in (usually water-based solvents), and provide long-lasting coloration upon deposition in the dermis (Piccinini et al., 2015; Bäumler, 2020), while colorants like dyes are less preferred as they are soluble and are typically associated with temporary vivid colors (Piccinini et al., 2015). In order to classify the colorants into ‘pigments’ and ‘dyes’, the chemical lists provided by the books titled ‘Industrial Organic Pigments’ (Hunger and Schmidt, 2018), ‘Industrial Inorganic Pigments’ (Buxbaum and Pfaff, 2005), ‘Industrial Dyes’ (Hunger, 2002), and the lists of banned and non-banned pigments in the EU, were relied upon. For colorants not present in these resources, the experimental water solubility information was obtained for chemicals from the REACH Dossiers and PubChem (Supplementary Information; Table S2). Further, a literature analysis was conducted, wherein the water solubility information was obtained from published evidence and available chemical safety data sheets (Supplementary Information; Table S2). For chemicals lacking any such experimental or regulatory information, their predicted water solubility information was obtained from CompTox Chemicals Dashboard and SwissADME (Daina et al., 2017) (Supplementary Information; Table S2). Notably, the predictions from the 2 resources were similar. Based on consensus, chemicals that were considered water soluble, i.e., having solubility values >1g/L, were considered as dyes and the rest were categorized as pigments (Supplementary Information; Table S2). Notably, 2 colorants, C.I. Pigment Green 13 (CAS:148092-61-9) and C.I. Natural Red 22 (CAS:98225-55-9), were not catalogued in any literature evidence, and predicted water solubility values were also not available, and were thus classified as ‘other colorant’ as they were not classifiable into pigments or dyes (Table S2).

For chemicals that could not be classified into categories such as colorants, auxiliaries, and preservatives, published evidence was relied upon to identify the role of such chemicals in tattoo inks (Supplementary Information; Table S2). It was observed that some chemicals are involved as intermediates in the production of colorants (Booth, 2000; Lim and Shin, 2015), while a few others are a result of chemical degradation of colorants (Sun et al., 2017; Negi et al., 2022). Therefore, such chemicals were categorized into ‘intermediates’ and ‘degradants’, respectively. It was also observed that some of the chemicals are contaminants resulting from the manufacturing processes and were therefore categorized as ‘contaminants’ (Lehner et al., 2014; Laux et al., 2016; Fels et al., 2023). Through this comprehensive exercise, the 364 tattoo ink chemicals were classified into 8 categories, namely, pigments, dyes, intermediates, degradants, auxiliaries, preservatives, contaminants, and other colorants (Table S2).

### 2.3. Identification of diseases associated with tattoo ink chemicals

The Comparative Toxicogenomics Database (https://ctdbase.org) is a comprehensive resource that curates published information on effects of environmental chemicals on health (Davis et al., 2025). Specifically, the database provides information on the diseases associated with chemicals, with diseases indexed using the Medical Subject Headings (MeSH) vocabulary, along with the associated published evidence. Additionally, it provides the genes shared between the chemical and the disease, inferred from the curated chemical-gene and gene-disease associations within CTD. An inference score is also provided to understand the significance of the overlap. This inference score helps rank the associated diseases to gain more insights from such associations (King et al., 2012). In this study, these chemical-disease associations from CTD were utilized to gain further insights into adverse effects associated with tattoo chemicals.

First, the chemical-disease associations were downloaded from CTD and filtered based on the CASRN to select the subset of tattoo ink chemicals. Next, associations that were marked as ‘marker/mechanism’ were selected to obtain direct evidence (Supplementary Information). Thereafter, to gain further confidence in the disease associations, only those diseases that contained overlapping gene information and associated inference scores were considered. Subsequently, a stringent criterion of a minimum of 5 overlapping genes, followed by an inference score cut-off of 20, was applied to obtain high confidence associations (Supplementary Information). Through this extensive filtration process, 346 chemical-disease associations were obtained that comprised 52 tattoo chemicals and 121 diseases (Table S3). Next, the associated diseases were manually mapped to the Disease Ontology (DO) database (https://disease-ontology.org/do), and corresponding disease categories were obtained to gain a broader understanding of the associated diseases (Supplementary Information). Among the 121 associated diseases, 15 were identified as symptoms, while 29 could not be mapped to any entries in the DO database and were therefore categorized as ‘unclassified’ (Table S3).

### 2.4. Curation of toxicological information from REACH Dossiers

The European chemicals agency (ECHA) provides a public database of chemicals that are registered in REACH (https://chem.echa.europa.eu/). In particular, various chemical information, including chemical properties and experimental toxicological data, have been submitted by the registrant into a dossier for each chemical. In this study, the experimental toxicological information pertaining to tattoo chemicals were manually obtained from the publicly available chemical dossiers accessible at: https://chem.echa.europa.eu/ (last accessed on 3 July 2025) (Supplementary Information).

First, the chemicals were identified through search based on their CASRNs. This revealed that 23 of the 364 chemicals are not present in this resource (Table S4). Dossier availability was checked for each chemical, where 115 chemicals had no dossiers and a further 15 chemicals had dossiers marked as inactive, indicating that the registrant had ceased manufacture or import of the substance, and were therefore not considered further in the analysis (Table S4). For the remaining 211 chemicals, the active dossiers with registration role as ‘Lead’ were selected to obtain corresponding toxicological information. Within the dossiers, it was observed that there were multiple sources of evidence for each toxicological endpoint, arising from different experimental or READ-across based studies. In order to obtain high confidence experimental data from these dossiers, the submissions with adequacy of study marked as ‘Key study’, type of information as ‘experimental study’, and a reliance score denoted by Klimisch score (Klimisch et al., 1997) of 1 or 2, were considered. It was observed that 16 of the 211 chemicals lacked any such studies across different toxicological endpoints (Table S4). For the remaining 195 chemicals the available experimental data, including endpoints such as skin or eye irritation/corrosion, skin sensitization, genotoxicity and associated adverse effects were obtained (Table S4).

### 2.5. Construction of stressor-centric adverse outcome pathway (sAOP) networks for tattoo ink chemicals

Adverse Outcome Pathways (AOPs) framework was envisioned to capture the mechanisms underlying stressor-induced toxicities and streamline their risk assessment (Ankley et al., 2010). According to this framework, the biological events, termed Key Events (KEs), underlying the toxicity mechanism are organized into a sequential chain of events, starting from Molecular Initiating Event (MIE), and terminating at Adverse Outcome (AO) (Ankley et al., 2010; Villeneuve et al., 2014a, 2014b). These events are connected by causal links called Key Event Relationships (KERs) (Ankley et al., 2010; Villeneuve et al., 2014a, 2014b). AOPs are developed to be stressor-agnostic and designed for specific adverse outcomes (Ankley et al., 2010; Villeneuve et al., 2014a, 2014b). The stressor-centric AOP network aggregates different AOPs specific to a stressor, thereby providing a holistic understanding of the diversity in toxicity mechanisms associated with that stressor, and eventually aid in their risk assessment (Aguayo-Orozco et al., 2019; Sahoo et al., 2024b, 2024a). AOP-Wiki is the largest repository of AOPs developed globally and is continuously updated, making it a living document. Therefore, based on our earlier studies (Sahoo et al., 2024b, 2024a), a systematic network-based filtration procedure was first leveraged to identify the complete and connected AOPs (Supplementary Information). In brief, the latest AOP-Wiki information was first downloaded, and the information within each AOP was systematically cleaned to identify 385 complete and connected AOPs, referred to as curated AOPs (Supplementary Information; Tables S5-S7). Next, the adverse biological effects data pertaining to the tattoo chemicals was obtained from ToxCast (Dix et al., 2007), CTD (Davis et al., 2025) (https://ctdbase.org), DEDuCT (Karthikeyan et al., 2021a, 2019) (https://cb.imsc.res.in/deduct/), NeurotoxKb (Ravichandran et al., 2021) (https://cb.imsc.res.in/neurotoxkb/), and REACH chemical dossiers (https://chem.echa.europa.eu/) (Table S8), and systematically mapped with the KEs present within AOP-Wiki. The detailed explanation of the mapping procedure is provided in the Supplementary Information.

Briefly, the prototypical stressor information available in the AOP-Wiki was first utilized to link tattoo ink chemicals to the corresponding AOPs and their constituent KEs. Next, the active ToxCast assay endpoints associated with tattoo ink chemicals were curated from ToxCast invitrodb v4.2 after removing assay endpoints identified as cytotoxicity-associated responses (Judson et al., 2016), and the corresponding assay targets were manually mapped to KEs in the AOP-Wiki to identify potential MIEs. High-confidence chemical-gene-phenotype-disease (CGPD) tetramers were constructed from CTD, and the associated diseases were manually mapped to KEs based on their underlying biological processes. The associated phenotypes were mapped to KEs by incorporating neighboring parent and child Gene Ontology (GO) terms to capture semantically related biological processes (Ashburner et al., 2000), followed by overlap with GO annotations associated with KE process identifiers in the AOP-Wiki and manual curation to retain biologically relevant matches. Next, chemical-associated endocrine-mediated endpoints from DEDuCT and neurotoxic endpoints from NeurotoxKb were manually mapped to KEs based on their underlying biological processes. Finally, experimental toxicity data from REACH dossiers were manually mapped to KEs based on the underlying biological processes associated with the reported adverse effects and genotoxic endpoints. Together, these approaches enabled the identification of chemical-KE associations across multiple complementary evidence sources.

Thereafter, such chemical-KE links were merged with the curated AOPs to obtain a stressor-centric AOP network linking 151 tattoo chemicals to 362 AOPs (Supplementary Information; Table S9). These stressor-AOP associations were further characterized by the metrics such as coverage score, a ratio of mapped events to the total events present within AOP, and level of relevance to understand the kind of associated events (Supplementary Information; Table S9) (Sahoo et al., 2024a). A higher coverage score indicates that the available evidence associated with a chemical spans a larger proportion of the toxicity pathway, while the level of relevance provides insight into whether the mapped evidence encompasses key mechanistic stages, including the MIE, intermediate KEs, and AO, thereby supporting the associated AOP as a plausible toxicity mechanism for the chemical.

It should be noted that these chemical-AOP associations were derived from diverse sources and represent a compilation of observed adverse effects mapped onto the AOP framework. Further experimental validation is necessary to establish definitive causal links between these tattoo ink chemicals and the associated adverse outcomes.

### 2.6. Identification of cytokine or chemokine receptors associated with tattoo ink chemicals

Skin is not only a physical barrier, but also serves as an active immunological organ facilitating constant communication between immune and non-immune cells (Nestle et al., 2009; Kabashima et al., 2019). To gain insight into the immunological consequences of exposure through tattoo ink, the potential interactions between the tattoo ink chemicals and immune receptors were explored. Based on an earlier study by Ravichandran *et al*. (Karthikeyan et al., 2021b), a tripartite network comprising chemicals, cytokine receptors and cytokines was constructed to explore potential immunomodulatory effects of tattoo chemicals.

First, the target genes of tattoo ink chemicals were obtained from two resources, CTD (Davis et al., 2025) (https://ctdbase.org) and active assay endpoints from ToxCast invitrodb v4.2 (Feshuk et al., 2023; U.S. EPA, 2024). CTD provides species-specific chemical-gene or chemical-protein interaction information curated from published evidence. Furthermore, CTD includes additional annotations describing the type of interaction between the chemical and gene. In this study, human-specific binary interactions between tattoo ink chemicals and genes or proteins with at least one associated evidence were selected for further analysis. The interaction types were also reviewed to exclude those annotated as ‘response to substance’, ‘chemical abundance’, ‘chemical synthesis’ and ‘cotreatment’, as these were not informative of direct interactions. The USEPA’s ToxCast invitrodb v4.2 (Feshuk et al., 2023; U.S. EPA, 2024) dataset provides bioactivity data for a wide range of environmental chemicals. Based on a previous study (Sahoo et al., 2024a), active chemical-gene interaction data were obtained after filtering out spurious results associated with cytotoxicity-related endpoints (Supplementary Information).

Next, the cytokines and chemokines receptor genes were compiled from Cameron *et al*. (Cameron and Kelvin, 2013), KEGG BRITE (Kanehisa et al., 2016) database, and Guide to Pharmacology (Harding et al., 2024) database. NCBI gene (https://www.ncbi.nlm.nih.gov/gene/) database was utilized to obtain corresponding Entrez gene identifiers, and standardize the obtained gene symbols, resulting in the identification of 177 unique receptors. Additionally, these resources provided information on the cytokines and chemokines known to interact with each receptor. Specifically, KEGG BRITE and the Guide to Pharmacology offered curated interaction annotations, such as activation or inhibition, between cytokines or chemokines and their receptors in humans.

### 2.7. Web interface and database management system

An online resource, Tattoo Ink Chemicals and associated Toxicities Knowledgebase (TICToK), was developed to provide access to the curated dataset generated in this study. Importantly, TICToK offers easy access to the compiled chemical information for the 364 tattoo ink chemicals, including their standardized identifiers, chemical structures, physicochemical properties, and molecular descriptors. Further, admetSAR (Yang et al., 2019), pkCSM (Pires et al., 2015), SwissADME (Daina et al., 2017) and the vNN server (Schyman et al., 2017) were used to predict ADMET properties for chemicals based on their structures. These properties included the log Kp value for skin permeation, fraction unbound in humans, subcellular localization, cytotoxicity, maximum recommended tolerated dose (MRTD), micronucleus and skin sensitization potential.

In addition, TICToK provides compiled information on the associated diseases, toxicological endpoint data from REACH Dossiers and ToxCast, and the linked AOPs identified through the construction of stressor-centric AOPs in this study. TICToK also provides information on the regulatory coverage and hazard classifications for tattoo ink chemicals. TICToK is accessible at: https://cb.imsc.res.in/tictok/.

TICToK stores the compiled data using the MariaDB relational database system (https://mariadb.org/) and accesses it through Structure Query Language (SQL) queries. Its web interface is developed using PHP (https://www.php.net/), combined with custom-built HTML, CSS, and JavaScript libraries such as jQuery (https://jquery.com/) and Bootstrap 5 (https://getbootstrap.com/docs/5.0/). For visualizing data, the platform employs interactive charts powered by Google Charts (https://developers.google.com/chart/). The entire system is hosted on an Apache webserver (https://www.apache.org) running on a Debian 9.4 Linux environment.

## 3. Results and Discussion

### 3.1. Identification and functional classification of 364 tattoo ink chemicals

In this study, 364 tattoo ink chemicals were compiled from multiple regulatory and scientific resources (Methods; Figure 1a; Table S1). Among the different lists, the EU JRC report (Piccinini et al., 2015) contributed the highest number of tattoo ink chemicals (242), of which 157 chemicals were not reported by other sources (Figure 1b). This was followed by the Australian NICNAS reports (National Industrial Chemicals Notification and Assessment Scheme (NICNAS) and Australian Government, Department of Health, 2018, 2017) which provided 27 unique chemicals (Figure 1b). In contrast, the CompTox Chemicals Dashboard and the NORMAN-SLE represented an identical dataset rather than independent compilations, as both were derived from the same primary regulatory source (Methods). Overall, 157 chemicals were found to be present in more than one list, indicating overlap across the compiled regulatory and scientific sources (Figure 1b).

**Figure 1:**
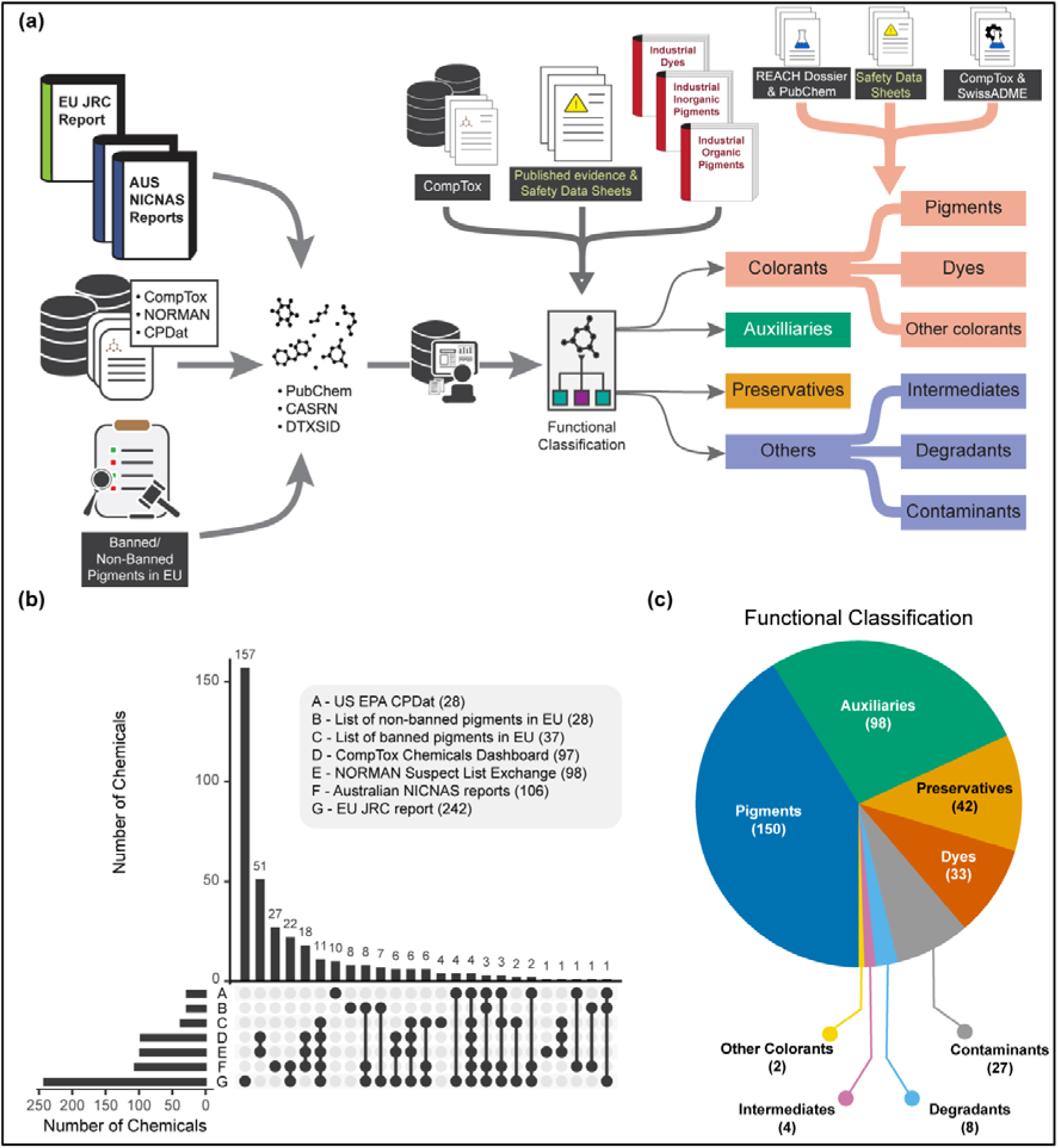
Identification of 364 unique tattoo ink chemicals from different sources. (a) The workflow to compile, curate and functionally classify tattoo ink chemicals. (b) UpSet plot illustrating the overlaps of tattoo ink chemicals across different sources. (c) Pie-chart depicting the distribution of functional types present in tattoo ink chemicals.

Next, based on the functions imparted by these chemicals in tattoo inks, 150 were categorized as pigments, 33 as dyes, 4 as intermediates, 8 as degradants, 98 as auxiliaries, 42 as preservatives, 27 as contaminants, and 2 as other colorants (Methods; Figure 1c; Table S2). It was observed that pigments account for the majority of the chemicals in our compilation, underscoring their central role in tattoo ink formulations as the primary agents responsible for imparting color to the dermal layers (Figure 1c). This was followed by a substantial number of auxiliaries, which are typically added to improve dispersion, stability, or performance of the pigments (Figure 1c) (Piccinini et al., 2015). Preservatives formed the next major category, while smaller numbers of dyes, other colorants, contaminants, intermediates, and degradants account for the remaining chemicals in our compilation (Figure 1c). It should be noted that these functional classifications were obtained directly from publicly available resources and used as reported for categorization.

### 3.2. Hazard classification reveals the presence of high-risk chemicals in tattoo inks

A better understanding of the nature of chemicals in tattoo inks will aid in directing further research and regulatory efforts. Therefore, the presence of carcinogens, skin sensitizers and irritants, endocrine disruptors, and neurotoxicants among tattoo ink chemicals was explored in this study based on existing resources (Figure 2a). These categories of chemicals are of particular concern as they have the potential to result in long-term effects (Grandjean and Landrigan, 2006; Diamanti-Kandarakis et al., 2009; Samet et al., 2020).

**Figure 2:**
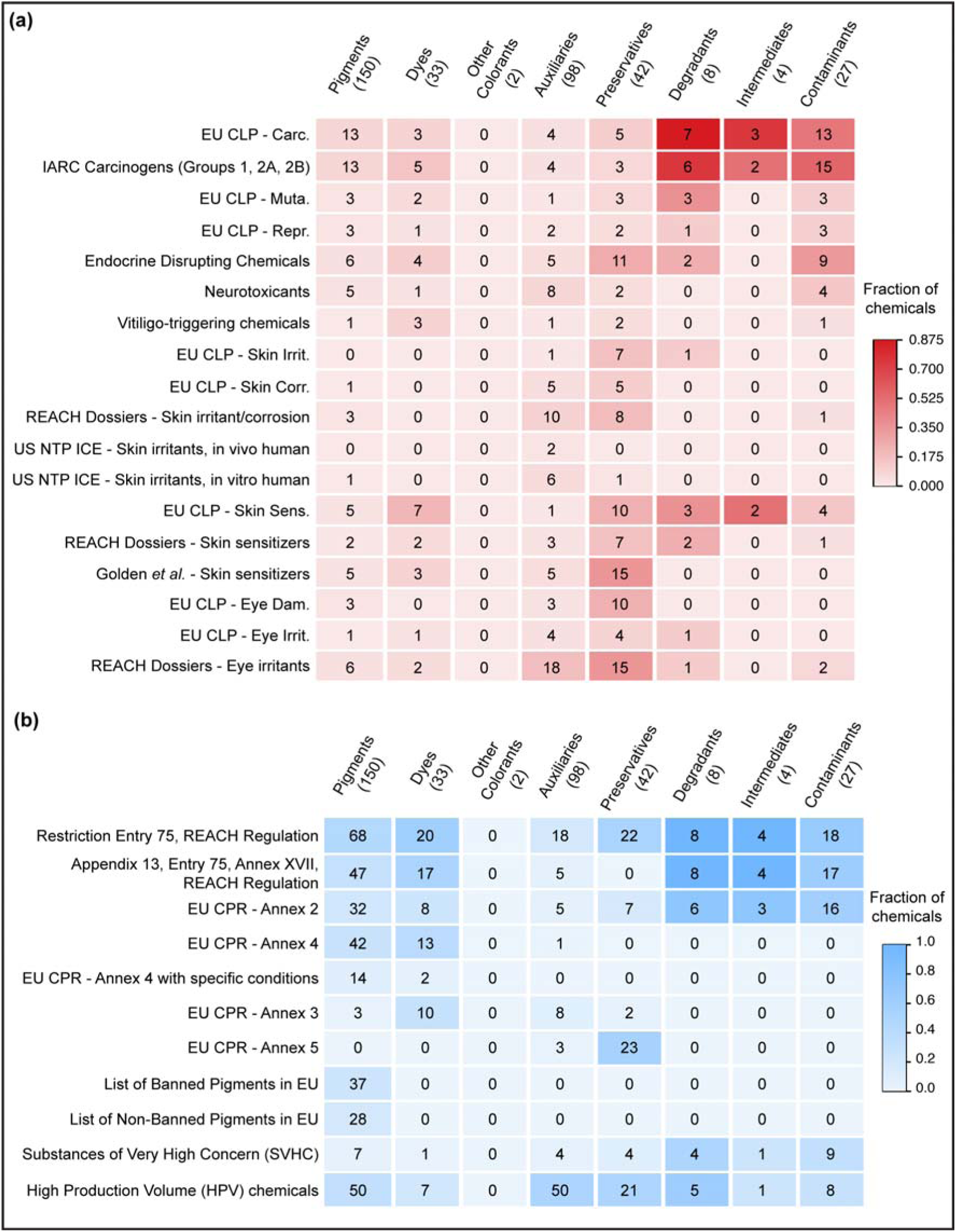
Hazard and regulatory characterization of tattoo ink chemicals. (a) Heatmap illustrating the presence of tattoo ink chemicals across various hazard resources. (b) Heatmap showing the presence of tattoo ink chemicals in existing regulations. In both heatmaps, the number of chemicals in each functional type and from different sources is provided.

The International Agency for Research on Cancer (IARC) provides monographs on the identification of carcinogenic hazards of chemicals to humans (https://monographs.iarc.who.int/list-of-classifications/; last accessed on 10 April 2025). According to the monograph, the chemicals are classified into 4 distinct categories: (i) Group 1: Carcinogenic to humans; (ii) Group 2A: Probably carcinogenic to humans; (iii) Group 2B: Possibly carcinogenic to humans; (iv) Group 3: Not classifiable as to their carcinogenic potential to humans. Among the tattoo chemicals, 10 are classified as Group 1, 5 classified as Group 2A, 33 classified as Group 2B, and 34 classified as Group 3 chemicals, based on the IARC monograph. In particular, the 48 tattoo chemicals classified into Groups 1, 2A, and 2B can be considered carcinogens (Figure 2a; Table S1) (Vincoff et al., 2024).

The harmonised classification and labelling (CLH) of hazardous substances under the Classification, Labelling and Packaging (CLP) Regulation (Regulation (EC) No 1272/2008) provide standardized labels to chemicals based on their associated hazards (European Parliament and the Council of the European Union, 2024). The CLH entries were obtained from ECHA chemicals database (https://chem.echa.europa.eu/obligation-lists/clhList; last accessed on 7 June 2026), and utilized to explore the hazards associated with 94 of the 364 tattoo chemicals in this study (Table S1). According to the classifications, carcinogens are categorized as follows (Terry et al., 2021): (i) Carc. 1A (H350): known human carcinogens; (ii) Carc. 1B (H350): presumed human carcinogens; (iii) Carc. 2 (H351): suspected human carcinogens. Among the 364 tattoo chemicals, it was observed that 4 are labelled as Carc. 1A, 32 as Carc. 1B, and 11 as Carc. 2 (Table S1). Notably, Gentian Violet (CAS:548-62-9) is labelled as Carc. 2, but in a mixture with ≥ 0.1% Michler’s ketone, it is labelled as Carc. 1B (Table S1). Comparison with the IARC classifications revealed that among the 48 chemicals considered carcinogens, two chemicals, indeno[1,2,3-cd]pyrene (CAS:193-39-5) and C.I. Direct Blue 15 (CAS:2429-74-5), which are classified as Group 2B chemicals, do not contain any CLH entries (Table S1), and are not covered in current tattoo-specific regulatory frameworks within the EU, highlighting gaps in the tattoo ink regulations in the EU. Furthermore, Acid Red 26 (CAS:3761-53-3) and C.I. Solvent Yellow 2 (CAS:60-11-7), which are classified as IARC Group 2B chemicals, are restricted for use in tattoo inks according to Appendix 13 to Restriction Entry 75 of Annex XVII to the REACH Regulation as amended by the Commission Regulation (EU) 2020/2081 (European Parliament and European Council, 2020), but do not contain CLH entries for carcinogenic hazards. Overall, integrating evidence from CLH entries and IARC monographs, 58 tattoo chemicals have been flagged for their carcinogenic potential (Table S1).

The Integrated Chemical Environment (ICE) (https://ice.ntp.niehs.nih.gov/) of the United States National Toxicology Program (US NTP) provides curated information on the *in vivo* and *in vitro* toxicological effects of chemicals, supporting hazard identification and risk assessment. Based on the skin irritation or corrosion data from ICE, 2 tattoo chemicals were identified as irritants/possible corrosives with *in vivo* human evidence (Figure 2a; Table S1), among which sodium hydroxide (CAS:1310-73-2) has CLH entry as Skin Corr. 1A (H314), while lactic acid (CAS:50-21-5) does not contain any CLH entry or is covered under any EU regulations for tattoo ink chemicals (Table S1). Based on the human *in vitro* evidence, 8 tattoo chemicals were identified as active skin irritants/corrosives of which 2 chemicals had *in vivo* evidence, while 4 others were identified as inactive (Figure 2a; Table S1). Among the 8 chemicals with *in vitro* evidence, 4 chemicals did not contain the CLH entry for skin corrosion (Table S1). Golden *et al*. (Golden et al., 2023) compiled a comprehensive list of skin sensitizers from various resources and categorized the chemicals as either active or inactive skin sensitizers. Based on this information, 28 of the 364 tattoo chemicals were identified as active skin sensitizers (Figure 2a; Table S1), of which 11 chemicals also contained CLH entry for skin sensitization, while 20 were inactive, of which 4-chloro-3,5-dimethylphenol (CAS:88-04-0) and 2-methyl-1,4-benzenediamine (CAS:95-70-5) contained CLH entry as Skin Sens. 1 (H317) (Table S1). The CLH entries also provide labels for skin sensitizers (Skin Sens. - H317), skin irritants (Skin Irrit. - H315), and skin corrosives (Skin Corr. - H314). Among the 364 tattoo chemicals, it was observed that 32 are skin sensitizers, 9 are skin irritants, and 11 are skin corrosives based on their respective CLH entries (Figure 2a; Table S1). In addition, CLH entries provide labels for eye damage (Eye Dam. - H318) and eye irritation (Eye Irrit. - H319). Among the 364 tattoo ink chemicals, 16 are classified as eye damage, and 11 are classified as eye irritants (Figure 2A; Table S1).

The REACH Dossiers (https://chem.echa.europa.eu/), maintained by the European Chemicals Agency (ECHA), provide comprehensive substance-level information, including toxicological profiles and hazard classifications for chemicals registered for use in the European Union. Based on the experimental information obtained from these dossiers, 22 tattoo chemicals were classified as active skin irritants/corrosives, 17 as active skin sensitizers, and 44 as active eye irritants (Methods; Figure 2a; Table S1). Among the 22 chemicals reported as skin irritants/corrosives, 11 chemicals do not contain any CLH entry for skin corrosion (Table S1). Furthermore, among the 17 skins sensitizers, 3 did not contain CLH entries for skin sensitization, of which 2,4,7,9-tetramethyl-5-decyne-4,7-diol (CAS:126-86-3) and cocamidopropyl betaine (CAS:61789-40-0) are not covered under any of the EU regulations for tattoo inks (Table S1). Among the 44 eye irritants, 25 chemicals do not contain CLH entries for eye irritation or damage, of which 12 chemicals are not covered under EU regulations for tattoo inks (Table S1). Overall, integrating the information from ICE, CLH entries, REACH dossiers and Golden *et al*. (Golden et al., 2023) skin sensitizers list, 29 active skin irritants/corrosives and 52 active skin sensitizers were identified among the 364 tattoo chemicals.

Additionally, 12 of 364 tattoo chemicals were classified as Repr. (H360/H361) in their CLH entries, and were thus identified as reproductive toxicants, and 15 chemicals were classified as Muta. (H340/H341) in their CLH entries, and identified as germ cell mutagens (Table S1). Notably, all these chemicals are already covered under the EU regulations for tattoo inks (Table S1). To further characterize the hazard potential of the identified tattoo chemicals, we cross-referenced them against manually curated literature-based knowledge resources including DEDuCT (Karthikeyan et al., 2021a, 2019) (https://cb.imsc.res.in/deduct/), NeurotoxKb (Ravichandran et al., 2021) (https://cb.imsc.res.in/neurotoxkb/), and ViCEKb (Chivukula et al., 2024) (https://cb.imsc.res.in/vicekb/). These resources compile experimental and clinical evidence from published literature to catalogue chemicals as potential endocrine disruptors, neurotoxicants, and vitiligo triggers, respectively. Among the 364 tattoo chemicals, 37 chemicals were identified as potential endocrine disrupting chemicals (EDCs) based on experimental evidence curated in DEDuCT, and 20 chemicals were identified as neurotoxicants based on their presence in NeurotoxKb (Figure 2a; Table S1). Among these, 9 potential EDCs and 3 neurotoxiciants are currently not covered in EU regulations for tattoo inks (Table S1). Additionally, 7 tattoo chemicals were catalogued as potential triggers of vitiligo in ViCEKb with ‘Patch test evidence’, and 1 tattoo chemical with clinical observation evidence (Figure 2a; Table S1). Notably, all these chemicals are already covered in EU regulations for tattoo inks (Table S1).

Importantly, no hierarchy or ranking has been applied across these different hazard resources, and all hazard classifications have been made available as independent sources of evidence (Table S1). This strategy facilitates Integrated Approaches to Testing and Assessment (IATA) by enabling researchers and regulatory assessors to integrate diverse evidence streams and apply expert judgment according to the context of the assessment (Caloni et al., 2022).

### 3.3. Existing regulatory coverage observed for tattoo ink chemicals

The primary EU regulatory instrument governing chemicals in tattoo inks is Restriction Entry 75 of Annex XVII to the REACH regulation (as amended by Commission Regulation (EU) 2020/2081) (European Parliament and European Council, 2020). According to this regulation, tattoo inks cannot be placed on the market in the EU if their constituent chemicals meet the following criteria:

- CLH entry as provided by Annex VI to Regulation (EC) no 1272/2008 (European Parliament and the Council of the European Union, 2024) is one of the following:

`o` Carc. 1A, 1B (H350) or 2 (H351), excluding those classified due to effects only following exposure by inhalation, and the concentration is ≥ 0.00005% by weight
`o` Muta. 1A, 1B (H340) or 2 (H341), excluding those classified due to effects only following exposure by inhalation, and the concentration is ≥ 0.00005% by weight
`o` Repr. 1A, 1B (H360) or 2 (H361), excluding those classified due to effects only following exposure by inhalation, and the concentration is ≥ 0.001% by weight
`o` Skin Sens. 1, 1A or 1B (H317), and the concentration is ≥ 0.001% by weight
`o` Skin Corr. 1, 1A, 1B or 1C (H314), and the concentration is ≥ 0.1% by weight when used as pH regulator or ≥ 0.01% by weight otherwise
`o` Skin Irrit. 2 (H315), and the concentration is ≥ 0.1% by weight when used as pH regulator or ≥ 0.01% by weight otherwise
`o` Eye Dam. 1 (H318), and the concentration is ≥ 0.1% by weight when used as pH regulator or ≥ 0.01% by weight otherwise
`o` Eye Irrit. 2 (H319), and the concentration is ≥ 0.1% by weight when used as pH regulator or ≥ 0.01% by weight otherwise
- Listed in Annex II to Regulation (EC) No 1223/2009, and the concentration is ≥ 0.00005% by weight
- Listed in Annex IV to Regulation (EC) No 1223/2009, with conditions specified in at least one of the columns g,h or i of the table in that Annex, and the concentration is ≥ 0.00005% by weight
- Listed in Appendix 13 to Entry 75 of Annex XVII to the REACH Regulation and exceeds the concentration limits specified in that Appendix

The CLH-based criteria listed above has been discussed in detail in the previous section. Regulation (EC) No 1223/2009 corresponds to the cosmetic products regulation (CPR) within the EU (European Parliament and the Council of the European Union, 2009), under which Annex II provides the list of substances prohibited for use in cosmetics products marketed in the EU, and Annex IV provides the list of allowed colorants in cosmetic products marketed in the EU. The chemical lists associated with the Annexes were accessed from CosIng, the official online database for cosmetic ingredients in the EU (https://ec.europa.eu/growth/tools-databases/cosing/reference/annexes; last accessed on 14 June 2025), and used to identify 77 tattoo chemicals in Annex II, and 56 in Annex IV, of which 16 chemicals have data in the specified columns according to the regulation (Figure 2b; Table S1). Furthermore, based on the chemicals listed in Appendix 13 to Restriction Entry 75 of Annex XVII to the REACH Regulation (European Parliament and European Council, 2020), it was observed that 98 of the 364 tattoo chemicals had specific concentration limits (Figure 2b; Table S1). Since the sources curated in this study do not contain corresponding concentration limits, 158 of 364 chemicals were identified as regulated tattoo chemicals in the EU based on satisfying criteria specified under Restriction Entry 75 of Annex XVII to the REACH Regulation, with the exception of the concentration limits (Figure 2b; Table S1).

To further explore existing regulatory coverage of tattoo chemicals, information provided by the other Annexes in CPR and ECHA were utilized. Based on the other annexes provided in the CPR, it was observed that 23 of 364 tattoo chemicals were present in Annex III that provides the list of substances that are restricted in cosmetic products marketed in the EU, and 26 were present in Annex V that provides the list of allowed preservatives in cosmetic products marketed in the EU (Figure 2b; Table S1) (European Parliament and the Council of the European Union, 2009). The inclusion of these annexes does not imply unrestricted permission for use in tattoo inks, and specific conditions defined in Restriction Entry 75 take precedence (Serup et al., 2025). Further, 30 of 364 tattoo chemicals were categorized as Substances of Very High Concern (SVHC) (https://echa.europa.eu/candidate-list-table), based on comprehensive evidence of carcinogenicity, reproductive toxicity, and endocrine-disrupting effects as evaluated under REACH (Figure 2b; Table S1). Moreover, 142 of 364 tattoo chemicals are listed as High Production Volume (HPV) (https://comptox.epa.gov/dashboard/chemical-lists/EPAHPV; https://hpvchemicals.oecd.org/ui/Search.aspx) chemicals, indicating their extensive industrial production (Figure 2b; Table S1). Among the 206 tattoo chemicals not currently covered in Restriction Entry 75, five chemicals, pyrene (CAS:129-00-0), benzo[g,h,i]perylene (CAS:191-24-2), fluoranthene (CAS:206-44-0), phenanthrene (CAS:85-01-8), and butylparaben (CAS:94-26-8), are cataloged as candidate chemicals in SVHC (Table S1), highlighting the potential risks associated with their presence in tattoo inks and PMU products. Of these five chemicals, the four polycyclic aromatic hydrocarbons are categorized as contaminants (Table S2). In contrast, butylparaben is categorized as a preservative (Table S2), indicating that it may be intentionally added to tattoo ink formulations. Therefore, further studies are needed to evaluate its toxicological effects in the context of tattoo ink use.

It should be noted that the regulatory mapping presented here represents an initial screening-level overview and is not a substitute for formal regulatory compliance assessment by qualified experts, and the regulatory status of these chemicals may change as new CLH entries are adopted or as Restriction Entry 75 is revised. Furthermore, the regulatory coverage presented in this study is based solely on the regulatory frameworks evaluated. Accordingly, the absence of a chemical from these regulations does not imply that it is safe or exempt from regulation, but only that it is not currently covered by the regulations examined in this study.

### 3.4. Chemical-disease associations reveal health risks linked to tattoo ink chemicals

To investigate the potential health impact of the tattoo ink chemicals, curated chemical–disease associations were extracted from the Comparative Toxicogenomics Database (CTD) (Methods). These associations provide evidence of previously reported adverse health effects linked to the chemicals of interest, including those used in tattoo inks. A stringent filtering criterion was applied to retain only the most significant chemical–disease associations, resulting in 121 diseases (spanning 19 disease categories including symptom and unclassified) that are associated with 52 chemicals, of which 11 are pigments, 4 are dyes, 1 degradant, 1 intermediate, 10 auxiliaries, 9 preservatives, and 16 contaminants (Methods; Figure 3, Table S3). Notably, cancer emerged as the most frequently associated disease class across all chemical categories (Figure 3).

**Figure 3:**
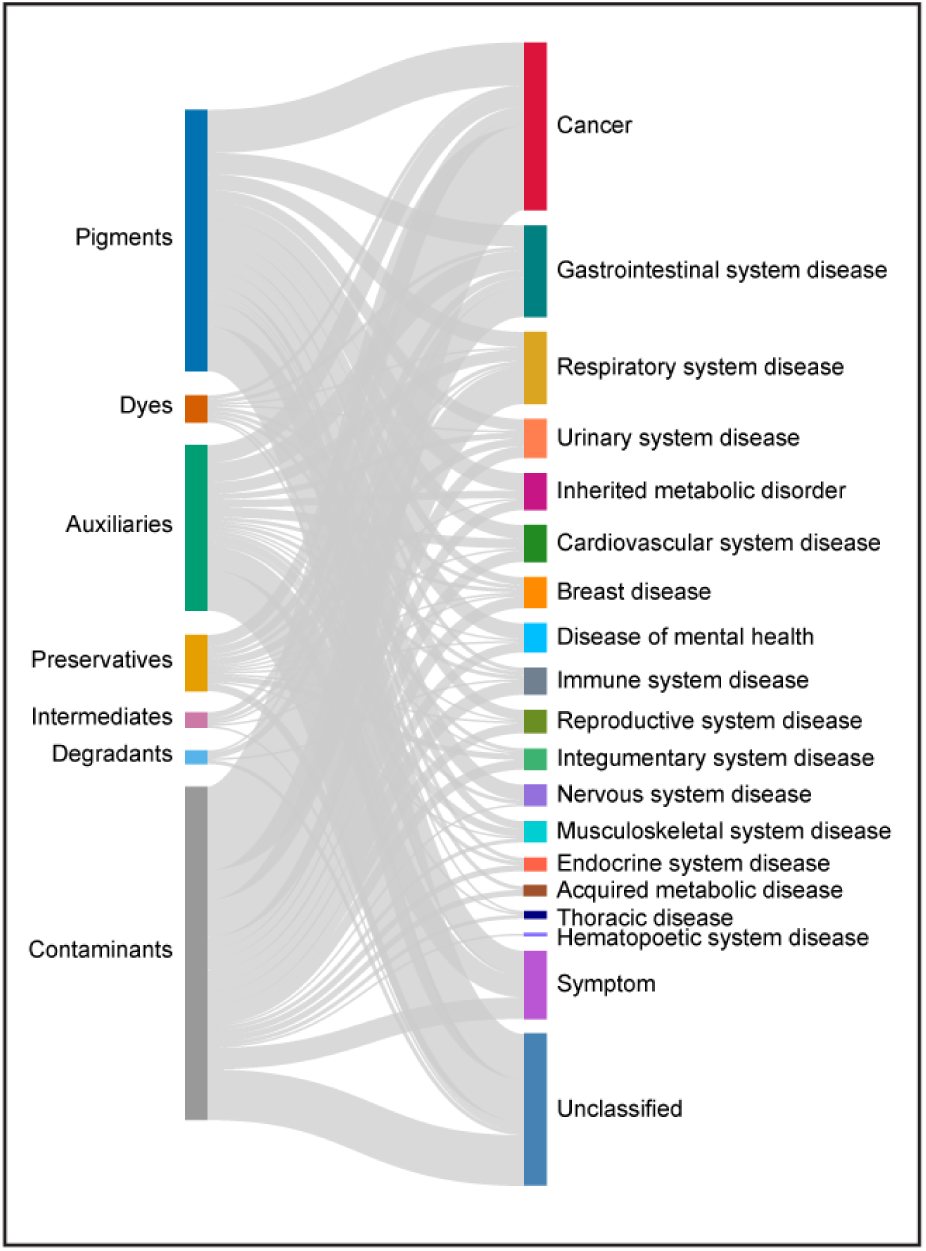
Aggregate visualization of the chemical-disease associations systematically compiled from Comparative Toxicogenomics Database (CTD). In this figure, the chemicals have been grouped based on their functional annotations, and diseases have been grouped based on their Disease Ontology (DO) classifications.

Contact dermatitis, including its immune-mediated form known as allergic contact dermatitis, is one of the most commonly reported skin conditions associated with exposure to tattoo inks and personal care products (Bassi et al., 2014; Shao et al., 2024). Through the CTD analysis, seven chemicals, namely 1,4-benzenediamine (CAS:106-50-3), phenol (CAS:108-95-2), 2-methyl-3(2H)-isothiazolone (CAS:2682-20-4), formaldehyde (CAS:50-00-0), nickel (CAS:7440-02-0), cobalt (CAS:7440-48-4) and dibutyl 1,2-benzenedicarboxylate (CAS:84-74-2), were associated with either contact dermatitis (MESH:D003877; DOID:2773), or allergic contact dermatitis (MESH:D017499; DOID:3042) or both (Table S4). Among these seven chemicals, six contained EU CLH entries for skin corrosion or skin sensitization, while 1,2-benzenedicarboxylate contained CLH entries for reproductive toxicity and acute aquatic toxicity (Table S1). Notably, all seven chemicals are currently regulated under the Restriction Entry 75 to REACH Regulation.

It should be noted that these chemical-disease associations were curated from published literature and are not specific to exposure through tattoo inks. Therefore, further experimental studies under conditions representative of tattoo ink exposure are required to evaluate the relevance of these associations in the context of tattooing.

### 3.5. Curated AOPs associated with tattoo chemicals

In this study, a data-centric approach was utilized, wherein the adverse events associated with the tattoo chemicals were systematically obtained from six resources, (Methods; Supplementary Information) and mapped to KEs within AOP-Wiki, resulting in 151 tattoo chemicals linked to 362 curated AOPs with varying coverage scores and levels of relevance (Methods; Table S9). Among these 362 AOPs 33 are endorsed, 18 are under review, and the rest are under development (Table S5). Figure 4a visualizes the stressor-AOP network of tattoo ink chemicals categorized as ‘pigments’, with level of relevance of 5 and coverage score cut-off of ≥ 0.4. In this network, the AOPs are classified into disease categories based on their adverse outcomes, and the stressor-AOP linkages are weighted based on their coverage scores (Methods).

**Figure 4:**
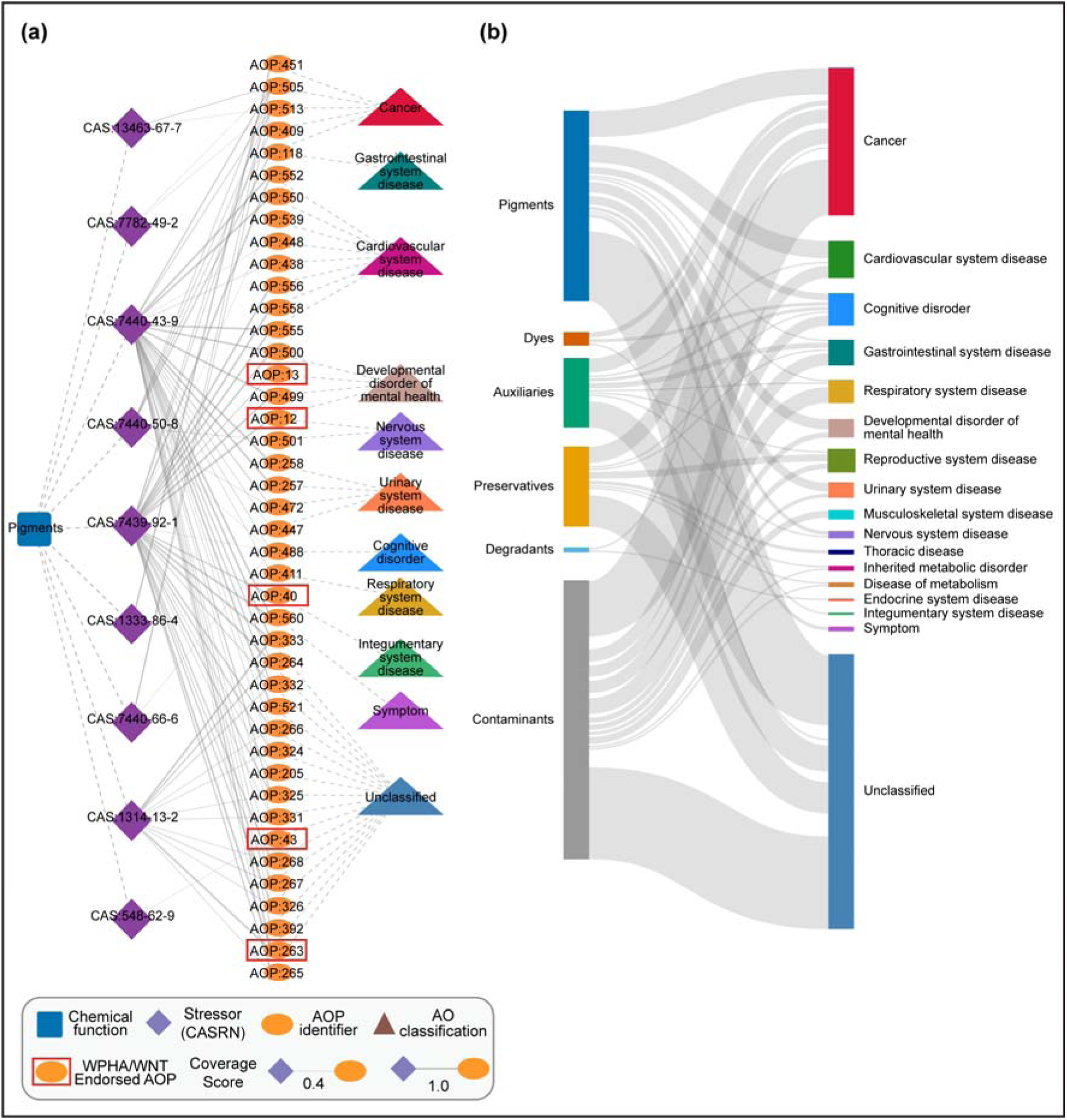
Stressor-AOP (sAOP) network for tattoo ink chemicals with coverage score ≥ 0.4, and level of relevance of 5. (a) Visualization of the sAOP network for tattoo ink chemicals classified as ‘Pigments’. The edges or stressor-AOP links are weighted based on their coverage score, i.e., the fraction of KEs within AOP that are linked with the pigments. (b) Aggregate visualization of the disease categories associated with AOPs in the sAOP network with level of relevance of 5 and coverage score ≥ 0.4 for each of the tattoo ink functional categories.

Further, 32 of these 151 chemicals were associated with 72 curated AOPs with level 5 relevance and a minimum coverage score of 0.4 (Figure 4b; Table S9). Notably, it was observed that 17 of these 32 tattoo chemicals are associated with AOP:505 (titled ‘Reactive Oxygen Species (ROS) formation leads to cancer via inflammation pathway’) and AOP:513 (titled ‘Reactive Oxygen (ROS) formation leads to cancer via Peroxisome proliferator-activated receptor (PPAR) pathway’), both of which are related with cancer. Although these AOPs are not endorsed by OECD, the weight of evidence supporting the pathway within these AOPs is ‘High’, and thus were utilized for further analysis (Supplementary Information; Table S5). Among the associated chemicals, 3 chemicals namely, ethanol (CAS:64-17-5), silica (CAS:7631-86-9), and butylparaben (CAS:94-26-8), have no limitations provided in Appendix 13 of Annex XVII under REACH Restriction Entry 75 for use in tattoo inks or EU CPR regulations for their use in cosmetic products. It should be noted that because the majority of these AOPs are not yet officially endorsed, they are utilized as plausible hypotheses to gain insights into chemical-induced toxicities, and may not be directly applicable for regulatory decision-making.

#### 3.5.1. Exploration of toxicity pathways associated with titanium dioxide

Titanium dioxide (TiO_2_; CAS:13463-67-7; CI 77891) is an inorganic pigment commonly used in tattoo inks, particularly to achieve white coloration (Piccinini et al., 2015). TiOc occurs naturally in three polymorphic forms, namely, rutile, anatase, and brookite (Wold, 1993). Among these, rutile and anatase can be readily produced (Wold, 1993), and have been detected in tattoo inks, with rutile being the predominant crystal form (Schreiver et al., 2017; Colboc et al., 2022; BfR, 2024). Anatase has been reported to exhibit greater cytotoxicity compared to rutile, primarily due to its higher photocatalytic activity (Schreiver et al., 2017; Colboc et al., 2022).

The hazard classification of TiO_2_ depends on the route of exposure, and should therefore be interpreted with caution when considering its regulatory status in the context of tattoo inks. The classification of TiO_2_ as IARC Group 2B (Table S1) and the earlier CLH entry as Category 2 carcinogen (Carc. 2, H350i) (European Commission, 2019), were based on evidence of lung tumour induction in rats following inhalation exposure to high concentrations of fine or ultrafine particles (Dankovic et al., 2007; BfR, 2024). Notably, this CLH entry is no longer valid as the Court of Justice of the European Union annulled the classification based on methodological grounds, and the European Court of Justice upheld this decision on 1 August 2025 (BfR, 2025). Therefore, there is currently no valid carcinogen classification that is applicable to intradermal exposure of TiO_2_ through tattooing, and it remains listed as an allowed colorant under EU cosmetics-relevant Regulation (EC) No 1223/2009, Annex IV (BfR, 2024; European Parliament and the Council of the European Union, 2009). Despite being widely regarded as safe in topical applications such as sunscreens and cosmetics (Dréno et al., 2019), the intradermal application of TiO_2_ via tattooing presents a different toxicological context, where direct deposition into the dermis raises concerns about persistent exposure and potential long-term effects (Skocaj et al., 2011; Schreiver et al., 2017).

To explore potential toxicity mechanisms underlying TiO_2_ exposure, the AOP framework was utilized. From the constructed stressor-AOP network, TiO_2_ was found to be associated with AOP:505 titled ‘Reactive Oxygen Species (ROS) formation leads to cancer via inflammation pathway’, and AOP:513 titled ‘Reactive Oxygen (ROS) formation leads to cancer via Peroxisome proliferation-activated receptor (PPAR) pathway’, both with a level of relevance of 5 and a coverage score of at least 0.4. These AOPs originate from the key event ‘Increase, Reactive oxygen species’ (KE:1115) and culminate in ‘Increase, Cancer’ (KE:885) as the adverse outcome (Figure 5). Here, AI-based tools, AOP-helpFinder (Jornod et al., 2022; Jaylet et al., 2025) (https://aop-helpfinder.u-paris-sciences.fr/), and Abstract Sifter (Baker et al., 2017) (https://comptox.epa.gov/dashboard/chemical/pubmed-abstract-sifter/), were leveraged to identify skin-relevant experimental studies in humans, linking TiO_2_ to the various biological events in these pathways (Table S10). Based on the available evidence, AOP:505 was selected for further exploration of the potential toxicity mechanism associated with TiO_2_ exposure.

**Figure 5:**
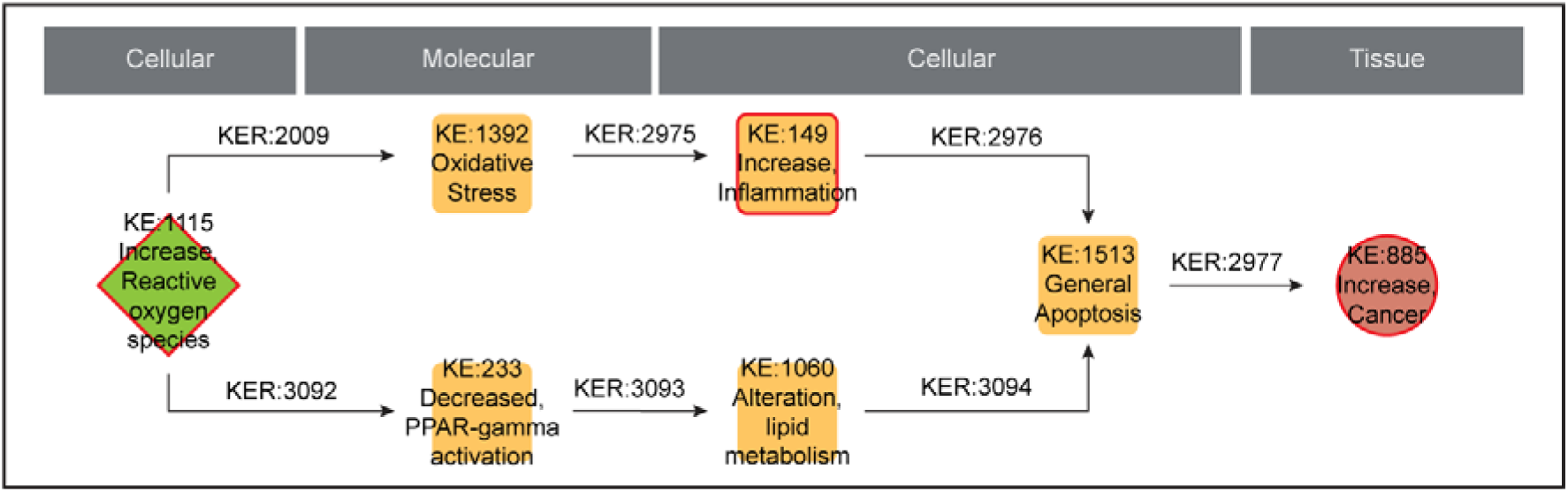
Toxicity pathways explored in the case study of titanium dioxide. The KEs with ‘red’ border were identified from the data integrative approach.

Different experiments have shown that exposure to both anatase and rutile forms of TiO_2_ leads to an increase in reactive oxygen species (ROS), and eventually, oxidative stress in both keratinocytes and epidermal cells (Xue et al., 2010; Shukla et al., 2011a, 2011b), consistent with MIE titled ‘Increase, Reactive oxygen species’ (KE:1115) and KE ‘Oxidative Stress’ (KE:1392). Subsequently, exposure to both anatase and rutile forms of TiO_2_ has also been observed to trigger inflammatory pathways in keratinocytes and Human Umbilical Vein Endothelial Cells (HUVECs), and potentially leads to apoptosis in these cells (Schanen et al., 2009; Shukla et al., 2011a, 2011b; Wright et al., 2017). TiO_2_ nanoparticle exposure has been observed to induce lung tumors in rats based on inhalation (Dankovic et al., 2007), potentially through prolonged inflammation, as indicated by a decrease in inhibitors of the NF-κB signaling pathway (Liu et al., 2017). Although inhalation is not directly relevant to tattoo ink exposure, TiO_2_ exposure has also been reported to upregulate pro-inflammatory cytokine secretion in HUVECs (Schanen et al., 2009), suggesting activation of the NF-κB signaling pathway (Lawrence, 2009; Schanen et al., 2009). These observations suggest that despite differences in route of exposure, activation of NF-κB signaling pathway represents a common downstream inflammatory response to TiO_2_ exposure. Since NF-κB pathway is a known contributor to tumor-promoting inflammation (Lawrence, 2009; Xia et al., 2014), this pathway may potentially link TiO_2_ exposure to cancer development in humans (Amiri and Richmond, 2005; Ueda and Richmond, 2006). Thus, by leveraging published evidence, we were able to explore the potential toxicity mechanism through which TiO_2_ may contribute to cancer development in humans.

It should be noted that this case study serves to evaluate mechanistic hypotheses derived from the AOP framework. Therefore, further experimental validation under conditions representative of intradermal exposure to TiO_2_ containing tattoo inks is needed to establish these potential adverse effects.

### 3.6. Systems biology approach uncovers potential immunomodulatory effects of tattoo ink chemicals

Cytokines and chemokines are signaling proteins that regulate various processes in the skin, including inflammation, wound healing, and tissue remodeling, in response to environmental chemical exposures (Coondoo, 2011; Lin et al., 2023). In this study, a systems biology approach was utilized to gain insight into the potential effects of tattoo ink chemicals on cytokine signaling pathways. First, the target genes or proteins for 156 tattoo ink chemicals were obtained from ToxCast invitrodb v4.2 and CTD (Methods). These targets were then analyzed to identify 40 chemicals interacting with 100 cytokine and chemokine receptors, which in turn interact with 105 distinct cytokines and chemokines (Table S11), highlighting a wide range of potential immune perturbations. Among the 40 chemicals, it was observed that 10 are pigments, 2 are dyes, 5 are auxiliaries, 12 are preservatives, and 11 are contaminants. The interactions for the 2 dyes are visualized as a tripartite network linking tattoo chemicals, receptors, and their associated cytokines or chemokines (Figure S1).

ToxCast screening assay data revealed that 3-iodo-2-propynyl-N-butylcarbamate (CAS:55406-53-6), C.I. Solvent Yellow 1 (CAS:60-09-3), and o-aminoazotoluene (CAS:97-56-3) activate the FAS receptor, while Carminic acid (CAS:1260-17-9) activates the CD40 receptor (Table S11). Additionally, 10 tattoo chemicals were found to inhibit key immune receptors such as IL6R, TNFRSF1A, and CD40 (Table S11). Among the tattoo chemicals analyzed, nickel (CAS:7440-02-0) was found to interact with the highest number of receptors, while the FAS receptor was targeted by 17 chemicals (Table S11). Notably, nickel can be introduced into the skin not only through ink formulations but also via abrasive wear of tattoo needles during application, underscoring the potential for widespread nickel exposure even when not explicitly listed as an ink component (Schreiver et al., 2019). The FAS receptor plays an important role in maintaining immune homeostasis by inducing apoptosis in activated or damaged cells in the skin (Strasser et al., 2009). Dysregulation of FAS-mediated signaling has been linked to autoimmune skin diseases, inflammatory responses, and cancer progression, suggesting that its perturbation by tattoo chemicals may contribute to long-term dermal diseases (Nagata and Golstein, 1995; Hill et al., 1999). Moreover, preservatives such as propylparaben (CAS:94-13-3) has been documented as skin irritant based on *in vivo* experiments conducted by US NTP (Table S1). Additionally, preservatives such as 1,2-Dibromo-2,4-dicyanobutane (CAS:35691-65-7), and auxiliaries such as lactic acid (CAS:50-21-5) have been documented as active skin sensitizers in list compiled by Golden *et al*. (Golden et al., 2023) (Table S1). However, these chemicals have not been restricted for use in tattoo inks based on the regulatory frameworks evaluated in this study (Table S1).

It should be noted that this is an exploratory analysis based on chemical-target interaction data obtained from existing literature curated in CTD and *in vitro* experimental observations available through ToxCast. CTD-derived chemical-gene interactions are subject to publication bias, resulting in the overrepresentation of well-studied chemicals. Therefore, the absence of a recorded interaction should be interpreted as a data gap rather than evidence of biological inactivity. The ToxCast assay data represent high-throughput *in vitro* evidence of chemical-target interactions and should be interpreted with caution, as these interactions may not directly translate to *in vivo* biological responses. Therefore, these interactions should be considered potential immunomodulatory effects of tattoo ink chemicals, and further experimental validation is needed to confirm these biological effects in tattooed skin.

### 3.7. TICToK: Tattoo Ink Chemicals and associated Toxicities Knowledgebase

In this study, a comprehensive list of tattoo ink chemicals was first compiled and curated. Then, various hazards associated with these chemicals were identified using multiple regulatory and scientific resources. Existing regulations were cross-referenced to identify existing regulatory coverage of these tattoo ink chemicals. Furthermore, toxicological endpoints associated with these chemicals were systematically curated and harmonized within the Adverse Outcome Pathway (AOP) framework. These associations provide potential toxicity mechanisms associated with the tattoo chemicals that require further experimental validation before they can support their regulatory assessment. All findings and generated results are made accessible through an online database, Tattoo Ink Chemicals and associated Toxicities Knowledgebase (TICToK), which is accessible at: https://cb.imsc.res.in/tictok.

The TICToK web interface allows user-friendly navigation of the database through its BROWSE and SEARCH functions (Figures S2-S3). Tattoo ink chemicals can be browsed based on their functional categories such as pigments, dyes, etc. An ADVANCED SEARCH option is available to query chemicals based on structural or physicochemical similarity (Figure S3). For each chemical, TICToK provides access to a wide range of chemical information including molecular structures, physicochemical properties, predicted ADMET values, and structural descriptors computed using PaDEL software (Figure S4). Additionally, TICToK displays toxicological endpoints compiled from ToxCast and REACH Dossiers, along with the associated AOPs identified in this study (Figure S4). All datasets compiled within TICToK are available as flat files in the DOWNLOAD section. TICToK also includes a HELP section, which provides a glossary and tutorials for navigating and using the online database.

TICToK provides standardized identifiers, comprehensive metadata, and downloadable datasets, ensuring findability, accessibility, interoperability and reusability, in compliance with FAIR principles (Wilkinson et al., 2016), and promotes transparency and sustainability in compliance with TRUST principles (Lin et al., 2020). Environmental scientists and regulators can utilize TICToK to identify tattoo ink chemicals that are also detected in environmental samples, enabling cross-referencing of hazard data, informing broader environmental risk assessments, and guiding the development of green chemistry alternatives for tattoo ink formulations.

#### 3.7.1. Utility of TICToK for investigating individual tattoo ink chemicals

To illustrate the utility of the TICToK webserver, zinc oxide was selected as a representative tattoo ink ingredient. Zinc oxide has been documented as a white pigment in tattoo ink formulations in the EU Joint Research Centre (JRC) report (Table S1), and is listed as a permitted colorant in cosmetics in the EU based on CPR (Table S1). However, the introduction of zinc oxide through tattooing and its persistence within the skin present a different toxicological context. TICToK was therefore examined to explore the evidence available on the hazards of zinc oxide in the context of tattooing.

Using the webserver’s search interface, zinc oxide was queried and its dedicated information page was accessed. This page provides key chemical identifiers, structural information and links to external resources such as CAS and the CompTox Chemicals Dashboard, together with physicochemical properties, predicted ADMET properties and descriptors relevant to computational analyses. The ‘Associated Hazards’ Tab showed that zinc oxide has two CLH entries, namely Aquatic Acute and Aquatic Chronic, indicating that available evidence concerns environmental hazards rather than human dermal endpoints. The ‘Regulatory Coverage’ tab listed zinc oxide in selected regulatory inventories, including as an allowed colorant for cosmetics, while no restriction is listed under Appendix 13 to Restriction Entry 75 of Annex XVII to the REACH Regulation. The Associated AOPs tab showed that zinc oxide is associated with 11 AOPs with a Level 5 relevance score. Among these, only AOP titled “Uncoupling of oxidative phosphorylation leading to growth inhibition via decreased cell proliferation” (AOP:263) is an endorsed AOP and is applicable to humans. The tab also provides the corresponding mapped key events and supporting biological evidence, enabling users to further examine the mechanistic evidence underlying these associations. The ‘ToxCast Endpoints’ tab did not identify any active ToxCast assays for zinc oxide following application of the activity and cytotoxicity filtering criteria, indicating the absence of high-confidence ToxCast evidence for this chemical in TICToK. Considered together with the evidence presented in the ‘Associated AOPs’ tab, these findings suggest that currently available mechanistic and *in vitro* evidence for zinc oxide is limited, particularly with respect to dermal toxicity following tattooing.

Overall, this example demonstrates how TICToK provides a single point of access to integrated chemical, physicochemical, toxicological, regulatory, and mechanistic information for individual tattoo ink chemicals. By allowing users to assess the available evidence across multiple domains, TICToK can help identify chemicals, such as zinc oxide, for which important data gaps remain and thereby support the prioritization of chemicals for targeted experimental or computational follow-up studies to evaluate potential skin-specific adverse effects.

### 3.8. Environmental effects associated with tattoo ink chemicals

While the presence of various hazardous chemicals in tattoo inks raises significant human health concerns, these chemicals may also pose ecological risks upon release into the environment. Many pigments and additives used in tattoo inks are not manufactured exclusively for tattoo applications but are also widely used across other industrial sectors, including textiles, paints, cosmetics, and advanced materials (Singh et al., 2025). Consequently, these chemicals may be released into the environment during different stages of their production, use, and disposal (Singh et al., 2025).

To specifically check for chemical exposure from tattooing, Kochs *et al*. (Kochs et al., 2025) conducted a comprehensive human exposure study to track the biokinetics and fate of soluble tattoo ink constituents following application. In particular, this study used hazard-free tracer molecules, namely potassium iodide, 4-aminobenzoic acid (PABA), and 2-phenoxyethanol (PEtOH), as safe substitutes and quantified the tracers and their metabolites in tattoo-associated consumables, blood, and urine samples (Kochs et al., 2025). Although the primary objective of this study was to estimate human exposure following tattooing, the quantification of tattoo ink tracers in tattoo-associated consumables and urine indicates that tattoo ink constituents may undergo multiple processes following application, including retention in consumables, systemic absorption, metabolism, and urinary excretion. Moreover, these observations may provide insight into potential pathways through which tattoo ink chemicals could subsequently enter the environment.

Once released into the environment, the behaviour of tattoo ink chemicals is expected to depend on their physicochemical properties. For instance, pigments that are generally insoluble are likely to persist in the environment, accumulate in sediments and organisms, and present long-term challenges for remediation and bioaccumulation (Takahashi et al., 2012; Pang et al., 2021). In contrast, dyes and auxiliaries, being more water soluble, can disperse more readily through aquatic systems, thereby increasing their environmental mobility (Mishra et al., 2020; Singh et al., 2024). Preservatives, due to their biocidal properties, may continue to impact microbial communities, potentially disrupting essential ecological functions (Hernández-Moreno et al., 2019; Reiß et al., 2021). Thus, the functional classification presented in this study may also offer valuable insight into the environmental fate and transport of tattoo ink chemicals, and could guide the development of safe and more sustainable alternatives for tattoo inks.

Overall, the available evidence offers a preliminary understanding of the potential environmental release and adverse effects associated with tattoo ink chemicals, and more targeted studies are required to establish their environmental hazards.

## 4. Conclusion

This study presents one of the most comprehensive efforts to date to compile, curate, and classify 364 chemicals present in tattoo inks and PMUs. It provides a systematic overview of the regulatory coverage and hazards associated with tattoo ink chemicals based on different EU regulations and scientific knowledgebases. Additionally, chemical-disease associations retrieved from CTD implicated a wide range of diseases, including skin conditions such as dermatitis that are already known to be associated with tattoo ink exposure. Stressor-AOP network analysis linked 151 tattoo chemicals to 362 curated AOPs from AOP-Wiki, while a systems biology approach revealed potential immunomodulatory effects through interactions with cytokine receptors. Together, these computational analyses explore mechanistic hypotheses linking tattoo ink chemicals to potential biological effects. All compiled chemical information, associated hazards, and regulatory coverage have been made available through the associated online database namely, TICToK accessible at https://cb.imsc.res.in/tictok.

Despite the broad scope of computational analysis in this study, certain limitations were recognized. The sources used to compile the list of tattoo ink chemicals did not specify the exact concentrations present in tattoo formulations, and as a result, concentration data were not captured in this study. While large-scale, publicly available chemical resources were used for the functional classification of tattoo ink chemicals, their inherent discrepancies may influence the accuracy of current classifications. Although tattoo inks are complex mixtures, the analysis focused on hazards associated with individual chemicals. Despite applying filters based on study type and reliance score to obtain toxicological data from REACH dossiers, an expert judgment is still required to resolve any inconsistencies in the data. While a systematic filter was applied to identify 385 curated AOPs from AOP-Wiki that were both connected and complete, only 35 of these were endorsed, 19 were under review, and the remainder were still in the development phase, indicating that many of these AOPs represent potential mechanistic hypotheses rather than established toxicological pathways. Additionally, the phenotype-based identification of associated KEs within the AOP-Wiki is constrained by the incomplete GO annotations. Although the current strategy broadly maps the biological events associated with chemical-induced adverse effects, it may miss specific toxicological contexts that require further experimentation. Furthermore, the potential long-term effects of degradation products from certain colorants were not explored due to limited experimental evidence on their breakdown mechanisms. The hazard classifications and regulatory coverage presented here reflect the status of the respective regulations and scientific resources at the time of data collection.

Nonetheless, this study is one of the most comprehensive efforts toward compiling and classifying chemicals present in tattoo inks, and assembling the associated hazards, regulations and toxicological information from diverse resources. The associated online database TICToK provides easy access to the chemical structures, physicochemical properties, molecular descriptors, and predicted ADMET properties, and will aid in future research towards tattoo chemical toxicology. Although not all associated AOPs are endorsed, many provide mechanistic hypotheses that were further supported with existing experimental evidence, offering insights into potential toxicity mechanisms and streamlining future research on these chemicals. Importantly, the study offers a broad regulatory mapping that highlights coverage of tattoo ink chemicals under existing EU regulations, including Restriction Entry 75 of the REACH Regulation, EU CPR and SVHC chemical lists. Future integration of data from published scientific literature could enrich the database, and remains a potential avenue for further development of this resource. Beyond human health, the functional classification presented here may also offer valuable insights into the environmental fate and transport of tattoo ink chemicals, as pigments may persist and accumulate in sediments and organisms, while dyes and auxiliaries may disperse through aquatic systems, and preservatives may disrupt microbial communities due to their biocidal properties (Takahashi et al., 2012; Hernández-Moreno et al., 2019; Mishra et al., 2020; Pang et al., 2021; Reiß et al., 2021; Singh et al., 2024). Altogether, this study is expected to serve as a valuable tool for researchers, regulators, and public health professionals interested in the safety evaluation of tattoo ink chemicals.

## Data availability

The data associated with this study is contained in the article or in the supplementary material, or in the associated online database accessible at https://cb.imsc.res.in/tictok/.

## Acknowledgements

Areejit Samal would like to acknowledge support from the Department of Atomic Energy (DAE), Government of India via Apex project to The Institute of Mathematical Sciences (IMSc) Chennai, and the ‘Marine Ecotoxicology and Ecological Risk Assessment’ (MEERA) Programme [MoES/OSMART/EFC/2021 dated 07.03.2022] at the National Centre for Coastal Research (NCCR), Ministry of Earth Sciences, Government of India. Vimal Kishore and Ajay Vikram Singh acknowledge the support received from Institute of Eminence (IoE), Banaras Hindu University, India. The authors thank Michael Giulbudagian and Peter Laux for their valuable suggestions and insightful comments that helped improve the quality of this work.

## CRediT author contribution statement

**Nikhil Chivukula:** Conceptualization, Data Compilation, Data Curation, Formal Analysis, Methodology, Software, Visualization, Writing; **Shreyes Rajan Madgaonkar**: Conceptualization, Data Compilation, Data Curation, Formal Analysis, Methodology, Visualization, Writing; **Shambanagouda Rudragouda Marigoudar:** Formal Analysis, Methodology, Writing; **Krishna Venkatarama Sharma:** Formal Analysis, Methodology, Writing; **Vimal Kishore:** Formal Analysis, Methodology, Writing; **Ajay Vikram Singh:** Conceptualization, Formal Analysis, Methodology, Writing; **Areejit Samal:** Conceptualization, Supervision, Formal Analysis, Methodology, Writing.

## Supplementary Figure Captions

**Figure S1:** Visualization of the tattoo ink chemical - cytokine/chemokine receptors - cytokines/chemokines tripartite network, comprising two tattoo ink chemicals classified as ‘Dyes’.

**Figure S2:** TICToK website. (a) The home page of TICToK web server with an easy navigation bar. (b) Browse page for chemicals based on their function in tattoo inks.

**Figure S3:** TICToK images of the ‘SEARCH’ sections. (a) The chemicals can be searched through their chemical identifiers from the top right corner of the page. (b) The search results in a tabulated page that contains different information pertaining to the search term. (c) ADVANCED SEARCH option based on physicochemical filtrations. (d) ADVANCED SEARCH option based on chemical similarity.

**Figure S4:** TICToK images of CHEMCIAL INFORMATION page. (a) Chemical identification page containing various chemical information and structure download files. (b) ‘Associated AOPs’ section containing information on the AOPs associated with the chemical. (c) ‘ToxCast Endpoints’ section contains the active toxicological endpoints associated with the chemical from ToxCast invitrodb v4.2.

## Supplementary Table Captions

**Table S1:** This table contains the chemical information for 364 unique tattoo ink chemicals identified in this study. For each chemical, this table provides the Chemical Abstracts Service Registry Number (CASRN), PubChem identifier, DSSTox identifier, chemical name, source(s) from which it was identified (separated by ‘|’), CASRN as provided in the sources, chemical/mixture/polymer categorization, SMILES, InChI, InChIKey, and ClassyFire classifications such as Kingdom, SuperClass, and Class. Additionally, for each chemical, the table provides the hazards, such as the International Agency for Research on Cancer (IARC) monographs on carcinogens to humans, DEDuCT endocrine disrupting chemical (EDC) class (https://cb.imsc.res.in/deduct/), neurotoxicants identified from NeurotoxKb (https://cb.imsc.res.in/neurotoxkb/), evidence associated with the vitiligo-triggering potential obtained from ViCEKb (https://cb.imsc.res.in/vicekb/), experimental evidence against chemical induced toxicities from United States (US) National Toxicology Program’s (NTP) Integrated Chemical Environment (ICE), active chemical information from REACH Dossiers experimental data (https://chem.echa.europa.eu/) on toxicity endpoints such as skin, eye irritants/corrosives, and skin sensitization, and Golden et al. (PMID:32388570) skin sensitizer classification. This table also provides different regulatory information pertaining to these chemicals such as High Production Volume chemicals (HPV) in the United States and Organisation for Economic Co-operation and Development (OECD) associated countries, Substances of Very High Concern (SVHC), list of banned and non-banned pigments in the EU, EU cosmetic products regulation (CPR) coverage (https://ec.europa.eu/growth/tools-databases/cosing/reference/annexes; last accessed on 14th June 2025), harmonised classification and labelling (CLH) under CLP regulations (Regulation (EC) No 1272/2008) (separated by ‘|’) as accessed from ECHA chemicals database (https://chem.echa.europa.eu/obligation-lists/clhList; last accessed on 7th June 2026), presence and corresponding weight restrictions in tattoos and permanent make-up based on Appendix 13 to the Restriction Entry 75 of Annex XVII of the REACH Regulation (Regulation (EC) No 1907/2006), as amended by Commission Regulation (EU) 2020/2081 (http://data.europa.eu/eli/reg/2020/2081/oj), and the chemicals regulated under the Restriction Entry 75.

**Table S2:** This table contains information on the functional classification of the 364 tattoo ink chemicals. For each chemical, the table provides the chemical function class, subclass, list source(s) from where the function was identified (separated by ‘|’), the literature source(s) from where the function was identified (separated by ‘|’), book source(s) from where the function was identified (separated by ‘|’), and water solubility information obtained from chemical safety data sheets, REACH Dossiers, PubChem, CompTox TEST predictions (in g/L), CompTox OPERA predictions (in g/L), and SwissADME prediction.

**Table S3:** This table contains the information on identified CTD chemical-disease associations for tattoo ink chemicals. For each chemical, the table provides the associated disease identifier and name, the literature source(s) (separated by ‘|’), the HGNC symbols of overlapping genes (separated by ‘|’), Inference score, and Disease Ontology information such as disease identifier, name and category (separated by ‘|’).

**Table S4:** This table contains the information on the toxicological endpoints, such as skin irritation/corrosion, eye irritation/corrosion, skin sensitization, repeat dose toxicity, genotoxicity, carcinogenicity, reproductive toxicity and developmental toxicity, associated with tattoo ink chemicals obtained from https://chem.echa.europa.eu/. Under each toxicological endpoint, the table provides the corresponding test guideline or experiment followed, the score based on Klimisch criteria, the corresponding result, and NOEC/NOAEL and/or LOEC/LOAEL values, wherever applicable.

**Table S5:** This table contains the curated list of 385 high confidence adverse outcome pathways from AOP-Wiki. For each AOP, the table provides the corresponding information on AOP identifier, AOP title, Handbook Version that was followed while construction of the AOP, Organisation for Economic Co-operation and Development (OECD) status, and taxonomic applicability of the AOP (separated by ‘|’ symbol). Additionally, for each AOP, the table provides the corresponding AOP identifier, computed fraction of KERs with ‘High’ evidence (i.e., F(High)), computed fraction of KERs with ‘Moderate’ evidence (i.e., F(Moderate)), computed fraction of KERs with ‘Low’ evidence (i.e., F(Low)), computed fraction of KERs with ‘Not Specified’ evidence (i.e., F(Not Specified)), computed cumulative weight of evidence (WoE).

**Table S6:** This table contains information on the 1228 Key Events (KEs) present in the curated list of 385 AOPs. For each KE, the table provides the corresponding information on KE identifier, KE title, level of biological organization, and associated AOP identifier(s) (separated by ‘|’ symbol).

**Table S7:** This table contains information on Key Event Relationships (KERs) present in each of the curated list of 385 AOPs. For each AOP, the table provides the AOP identifier, corresponding KER identifier, upstream KE identifier, downstream KE identifier, MIE(s) among upstream and downstream KEs (separated by ‘|’ symbol), AO(s) among upstream and downstream KEs (separated by ‘|’ symbol), adjacency of KER, weight of evidence of KER, and quantitative understanding of KER. Note that the KER identifiers starting with 10000 were manually assigned by the authors as the KER was mentioned in the AOP page but was not assigned an identifier in AOP-Wiki.

**Table S8:** This table contains information on 529 Key Events (KEs) from 385 high confidence AOPs that are associated with tattoo ink chemicals. For each KE, the table provides corresponding information on KE identifier and title, CAS chemical identifier (separated by ‘|’) for associated chemical(s), source(s) from which the associations are inferred (separated by ‘|’ symbol), AOP Identifier(s) from which KE mapping was inferred (separated by ‘|’ symbol), associated CTD disease(s) (separated by ‘|’ symbol), associated CTD phenotype(s) (separated by ‘|’ symbol), associated DEDuCT endpoint(s) (separated by ‘|’ symbol), associated NeurotoxKb endpoint(s) (separated by ‘|’ symbol), associated REACH Dossier experimental toxicological endpoints (seperated by ‘|’ symbol), and associated ToxCast assay endpoint(s) (represented as (assay_name:response))(separated by ‘|’ symbol).

**Table S9:** This table provides edge list for the complete stressor-AOP network for tattoo ink chemicals. For each edge in the network, the table provides the function associated with stressor, stressor, computed coverage score, level of relevance, AOP identifier, and the AO classification of the corresponding AOP.

**Table S10:** This table contains information on the literature evidence for the association of 7 KEs with titanium dioxide. These 7 KEs are present in the two AOPs linked to titanium dioxide in the stressor-AOP network based on ‘Level 5’ level of relevance and coverage score cut-off of 0.4. For each KE, the table provides the corresponding KE identifier, KE title, level of biological organization, KE type, organism in which titanium dioxide exposure is studied, study type, dosage information, the type of titanium dioxide tested, abbreviated description of association with titanium dioxide exposure, and the PubMed ID (PMID) of the corresponding reference.

**Table S11:** This table provides the information on the interactions of tattoo ink chemicals with cytokine/chemokine receptors, and their corresponding cytokine/chemokine interactions. For each chemical, the table provides, the source(s) from which the chemical-receptor interactions were obtained, the corresponding interaction action(s) (separated by ‘|’) and literature source(s) (separated by ‘|’) obtained from CTD, ToxCast assay endpoint name and corresponding response, and corresponding information on interacting receptors such as their NCBI gene ID, HGNC symbol, and source(s) (separated by ‘|’) from where it was identified. Additionally, for each of the receptors, the table provides information on the cytokine/chemokine-receptor interactions, such as the type of interactions and corresponding sources from which cytokine/chemokine-receptor interactions were identified (separated by ‘|’), and corresponding information on cytokines/chemokines such as their NCBI gene identifiers, HGNC symbol and source(s) from which they were identified (separated by ‘|’). In this table, the cytokines/chemokines and their receptors were identified from 3 sources, KEGG BRITE (KEGG), Cameron et al. (NCBK6294), and Guide to Pharmacology (GoP).

**Table S12:** This table contains information on 59342 chemical-gene-phenotype-disease (CGPD) tetramers constructed for tattoo ink chemicals from Comparative Toxicogenomics Database (CTD). For each tetramer, the table provides the corresponding information on the chemical CASRN, NCBI gene identifier, NCBI gene name, phenotype identifier, phenotype name, MESH disease identifier, and MESH disease name.

