## Supplementary material for "TICToK: A comprehensive knowledgebase of tattoo ink chemicals and investigation of their associated toxicities and regulations": Figure S

#### for

### Supplementary Text

#### 1. Identification of tattoo ink chemicals from different scientific and regulatory resources

In this study, a comprehensive list of chemicals in tattoo inks was systematically compiled from various resources. First, the list of chemicals in tattoo inks was downloaded from NORMAN Suspect List Exchange (NORMAN-SLE)[1] (https://www.norman-network.com/?q=suspect-list-exchange) (last accessed on 27 May 2025) list number S86 titles TATTOOINK, and CompTox Chemical Dashboard's[2] chemical list titled TATTOOINK (https://comptox.epa.gov/dashboard/chemical-lists/TATTOOINK) (last accessed on 27 May 2025). Both these lists are provided in structured formats and include various chemical information, such as their chemical names, Chemical Abstracts Service Registry Numbers (CASRN) (https://commonchemistry.cas.org/), PubChem chemical identifiers (https://pubchem.ncbi.nlm.nih.gov/), and Distributed Structure-Searchable Toxicity (DSSTox)[3] identifiers (DTXSID).

Next, the United States Chemical and Products Database (CPDat)[4] list of chemicals was accessed by downloading the 'Composition Data' and 'Function data' provided by the ChemExpo knowledgebase (https://comptox.epa.gov/chemexpo/get_data/) (last accessed on 27 May 2025). CPDat classifies the chemicals based on standardized product use category (PUC) terminology, wherein the tattoo ink chemicals are classified as 'tattoo inks' under the 'PUC Product Type' (https://comptox.epa.gov/chemexpo/puc/322/). CPDat further provides the cleaned CASRN for each chemical. The chemical functions associated with these tattoo ink chemicals were retrieved from the 'Function data' file, based on the CASRN of the chemicals.

Next, the European Union Joint Research Centre (EU JRC) report titled 'Safety of tattoos and permanent make-up'[5] was accessed to identify chemicals in tattoo inks. The report provides lists of various pigments, auxiliaries, and preservatives present in tattoo inks, in the chapter titled '4.3. Ingredients of tattoo and PMU inks and their fate'. For each chemical, the chapter also provides their CASRN and chemical names. These chemical lists were manually extracted from this chapter, and the CASRNs were manually reviewed and updated. Subsequently, the reports from the Australian Industrial Chemicals Introduction Scheme (formerly known as the National Industrial Chemicals Notification and Assessment Scheme (NICNAS)) titled 'Characterisation of tattoo inks used in Australia'[6] and 'Investigation of the composition and use of permanent make-up (PMU) inks in Australia',[7] provided lists of chemicals in their appendices. The chemical names and CASRN were manually extracted, reviewed, and updated.

Annex XVII of the REACH regulations provides the list of restricted chemicals in tattoo and PMU inks in the European Union (EU) (https://eur-lex.europa.eu/eli/reg/2020/2081/oj/eng). The pigments present in this annex were therefore compiled as a 'List of banned pigments in the EU'. Next, chemicals were compiled from the REACH-compliant safety data sheets of prominent tattoo ink suppliers operating in the EU and those not listed under Annex XVII of the REACH regulation were classified as not currently restricted. Thereafter, pigments were extracted from this list as 'List of non-banned pigments in the EU'.

The chemical names and corresponding CASRN were retrieved. Finally, the CASRNs were used to compile a unique list of 364 chemicals derived from these resources. Subsequently, chemical information, including chemical names, PubChem identifiers, and DSSTox identifiers, was manually gathered based on the CASRNs of the identified chemicals. Subsequently, two-dimensional (2D) structural data was retrieved from PubChem based on their compound identifiers (CIDs). For chemicals lacking defined structural information, chemical information within the PubChem was utilized to categorize the chemical as a mixture or polymer.

**2. Manual assignment of chemical functions within tattoo inks**

To classify colorants as either 'pigments' or 'dyes', chemical lists provided in the appendices of the books 'Industrial Organic Pigments'[8] and 'Industrial Dyes'[9] were consulted. Chemical Abstracts Service Registry Numbers (CASRNs) from the chemicals were matched against the identifiers extracted from these references to assign them as 'pigments' or 'dyes'. For inorganic pigments, CASRNs were individually searched within the book 'Industrial Inorganic Pigments',[10] and chemicals with matching identifiers were manually verified for their function as pigments.

For colorants that were still not classified, experimental water solubility information was sourced from various resources. Initially, water solubility data was obtained from REACH chemical dossiers and PubChem chemical pages. In REACH dossiers, the water solubility is measured according to guidelines outlined in OECD Test No. 105.[11] These guidelines set a threshold of 10 mg/L, with anything below considered insoluble. Chemicals tested experimentally were annotated as insoluble (<10 mg/L), slightly soluble (10-500 mg/L), moderately soluble (500-1000 mg/L), or soluble (>1g/L), depending on the solubility value. In addition, a reliability score, based on Klimisch score,[12] is also provided. Therefore, to gain proper experimental result, only chemicals for which the corresponding Klimisch score was 1 or 2 were selected. PubChem also offers water solubility information, including soluble, slightly soluble, and insoluble labels, which were used for verification. For chemicals lacking this information, chemical safety datasheets were checked, based on a Google search based on search query: chemical_name AND "safety data sheet", where chemical_name was replaced with that of chemical of interest. The results were manually screened to obtain relevant information. However, 10 chemicals still lacked experimental solubility data across all resources. For these chemicals, predicted water solubility was sourced from the CompTox Chemicals Dashboard[2] (https://comptox.epa.gov/dashboard/) and SwissADME.[13] SwissADME provides an interpretation of their predicted results, whereas CompTox offers predictions from TEST and OPERA models, with results in mols/L, which were converted to g/L by multiplying with the corresponding molecular weights, and interpreted using the criteria in REACH dossiers.

To identify chemicals as 'intermediates', 'degradants', or 'contaminants', searches were conducted on Google Scholar and Google using a structured search query. First, Google Scholar was searched for literature on the role of these chemicals using the following query: chemical_name AND "tattoo ink", where chemical_name was replaced with the name of the chemical. If this query did not return any results, Google was used to search for literature with the same query. Functional roles of chemicals in tattoo inks were identified by reviewing research articles and other literature sources retrieved through the query search, with only those sources containing relevant information included in the analysis. Additionally, to obtain relevant chemical safety data sheets, a Google search was constructed with the following structured search query: chemical_name AND "safety data sheet", where chemical_name was replaced with that of chemical of interest. The results were manually screened to obtain relevant information.

**3. Chemical-disease associations within Comprative Toxicogenomics Database**

Comparative Toxicogenomics Database (CTD) is the largest resource that systemically curated chemical-related information from published evidence, including disease associations. CTD annotates chemical-disease associations as 'markers/mechanism' if the chemical is correlated with or plays an important role in the associated disease ()https://ctdbase.org/help/glossary.jsp#cdactions). Additionally, CTD provides data on overlapping genes and an inference score to understand the significance of the overlap. This inference score helps rank the associated diseases to gain more insights from such associations.[14]

The associations were first filtered to identify those with the 'marker/mechanism' annotation, and contained at least one published evidence to support the association. Next, only associations with a minimum of five overlapping genes were considered, based on empirical findings from gene set enrichment analysis that suggested a minimum 5 gene overlap indicated a biological relevance over random event.[15] Furthermore, an inference score cut-off of 20 was applied, as it falls within the top quartile of all scores, ensuring that only the most significant associations were included in the analysis.

**4. Compilation of AOPs within AOP-Wiki**

Adverse Outcome Pathway (AOP) is a toxicological knowledge framework that captures biological processes underlying stressor-induced toxicity.[16] In this framework, toxicity-related biological events, referred to as Key Events (KEs), are organized sequentially, beginning with the interaction of the stressor with a biological target, known as the Molecular Initiating Event (MIE), and culminating in an Adverse Outcome (AO).[16–19] The causal, directional relationships between these events are termed Key Event Relationships (KERs).[16–19] AOPs are stressor-agnostic and context-specific, meaning each AOP is tailored to a particular biological scenario.[16–19] Developed globally, these AOPs are deposited in the AOP-Wiki (https://aopwiki.org), which is the largest, publicly accessible repository hosted by the Society for the Advancement of Adverse Outcome Pathways (SAAOP). AOP-Wiki hosts several AOPs, where each AOP is documented in the form of KEs (including MIE and AO) and KERs, all of which are supported by scientific evidence (https://aopwiki.org/handbooks/5). Therefore, in this study, AOP-Wiki was relied upon to obtain the latest available AOPs.

First, XML file (released 1 April 2025) was downloaded from the 'Project Downloads' page in AOP-Wiki to obtain the latest information on AOPs. Then, the data was parsed to extract information associated with AOPs like AOP identifier, AOP title, associated KEs (including MIEs and AOs) and KERs, linked stressors, handbook version followed for development of AOP, status according to OECD, and biological applicability information such as taxonomy, sex and life-stage of the organism, and their corresponding weight of evidence, using an in-house Python script. Additionally, for each KE, information like KE title, KE identifier, level of biological organization, action name, object name, object identifiers and process name, and information associated with KERs like upstream/downstream KEs, evidence for biological plausibility of KER, adjacency, and the extent of quantitative understanding of KER, were extracted.

**5. Identification of 'high confidence' AOPs within AOP-Wiki**

AOPs are considered living documents as they are collaboratively developed globally, and are continuously updated based on new scientific evidence (https://aopwiki.org/handbooks/5). Consequently, many AOPs may remain incomplete, or lack sufficient information for further analysis. Therefore, based on our previous studies,[20–22] each AOP was systematically checked to filter high quality and complete AOPs. First, AOPs were manually checked and those that comprised KEs with title as 'unknown', or lacked any KEs or KERs, were removed. Next, NetworkX library[23] was employed in Python to check for disconnected AOPs, and those that contained disconnected components were manually inspected and updated before filtration of AOPs with disconnected components. Finally, the presence of MIE, AO, and a directed path between them were checked, and those that lacked any, were filtered out. Through this combined manual and computational effort, 385 complete, connected and high quality AOPs were obtained from AOP-Wiki (last accessed on

18 June 2025), which were designated as 'curated AOPs' (Table S5). These 385 curated AOPs comprised 1228 unique KEs (Table S6) and 1966 unique KERs (Table S7).

**6. Computation of cumulative weight of evidence (WoE) of AOPs**

AOP-Wiki provides a weight of evidence (WoE) information for each KER in an AOP based on its biological plausibility (Table S7). The WoE is a qualitative score represented in terms of 'High', 'Moderate', 'Low' and 'Not Specified'. Following Ravichandran *et al.*,[24] the fraction of KERs within an AOP with 'High' WoE [represented as F(High)], 'Moderate' WoE [represented as F(Moderate)], 'Low' WoE [represented as F(Low)] and 'Not Specified' WoE [represented as F(Not Specified)], were computed and subsequently a cumulative WoE was assigned to the AOPs based on the following criteria:

i. If F(High) ≥ 0.5, the cumulative WoE of AOP is 'High'
ii. If F(High) < 0.5, but (F(High) + F(Moderate)) ≥ 0.5, the cumulative WoE of AOP is 'Moderate'
iii. If (F(High) + F(Moderate)) < 0.5, but (F(High) + F(Moderate) + F(Low)) ≥ 0.5, the cumulative WoE of AOP is 'Low'
iv. If none of the above criteria are satisfied, the cumulative WoE of AOP is 'Not Specified'

Table S5 provides the cumulative WoE of the curated AOPs.

**7. Identification of KEs associated with tattoo ink chemicals**

Among the goals of this study was to investigate the toxicities associated with tattoo ink chemicals using the AOP framework. Based on our previous studies,[20–22] different toxicogenomics and biological endpoint information related to tattoo ink chemicals was obtained from ToxCast,[25] Comparative Toxicogenomics Database (CTD)[26] (https://ctdbase.org), DEDuCT[27,28] (https://cb.imsc.res.in/deduct/), NeurotoxKb[29] (https://cb.imsc.res.in/neurotoxkb/), AOP-Wiki (https://aopwiki.org), and REACH Dossiers provided by European Chemical Agency (ECHA) (https://chem.echa.europa.eu/). The obtained data was then systematically integrated to identify KEs within AOP-Wiki that are linked to tattoo ink chemicals.

##### 7.1. Identification of KEs using ToxCast

ToxCast is a program by the United States Environmental Protection Agency (US EPA) designed to enhance chemical toxicity predictions through *in vitro* high-throughput screening of various environmental chemicals.[25] These *in vitro* approaches provide crucial information on the associated biological processes and gene alterations triggered by stressor interactions, which can aid in identification of potential MIEs related to active chemicals.[20–22,30–32] In this study, the latest version of ToxCast data, ToxCast invitrodb 4.2,[33,34] was utilized to identify KEs associated with tattoo ink chemicals.

First, the chemicals and their corresponding assay information were extracted from the 'mc5-6_winning_model_fits-flags_invitrodbv4_2_SEPT2024.csv' file, and active assay endpoints for each chemical (defined as 'hitc' ≥ 0.9) were identified.[34] Next, the 'top' value of the corresponding winning model from the 'mc4_all_model_fits_invitrodbv4_2_SEPT2024.Rdata' file was utilized to determine whether these active chemicals exhibited an 'activatory' or 'inhibitory' effect.[34]

To ensure that these active endpoints are not due to non-specific activation of reported genes (referred to as 'cytotoxicity-associated burst' phenomenon),[35] the following Z-score statistic proposed by Judson *et al.*[35] was applied:

$$Z(chemical, assay) = \frac{-logAC_{50}(chemical,assay) - median[-logAC_{50}(chemical,cytotox)]}{\text{global cytotoxicity MAD}}$$

Here, 'logAC$_{50}$(chemical,assay)' is the logarithm of the AC$_{50}$ value of the chemical in the assay, 'logAC$_{50}$(chemical,cytotox)' is the logarithm of the AC$_{50}$ value of the chemical in the corresponding cytotoxicity assay, and the 'global cytotoxicity MAD' is the median of the MAD (median absolute deviations) of the $\log AC_{50}$(chemical,cytotox) distributions across all chemicals. Based on this definition, Z-scores ranging between +3 and -3 are considered indicative of cytotoxicity-associated bursts.[35]

In this study, the global cytotoxicity MAD and $\log AC_{50}$(chemical,cytotox) (given by the column titled 'cytotox_median_log') were retrieved from the 'cytotox_invitrodb_v4_2_SEPT2024.xlsx' file to identify the cytotoxicity-associated bursts associated with tattoo ink chemicals.[34] The assays with Z-scores between +3 and -3 were discarded, and the resulting assays were subsequently mapped to KEs within AOP-Wiki.

The ToxCast assay endpoints, and the KE titles from the AOP-Wiki were manually inspected to identify relevant KEs. This manual curation led to the identification of 160 KEs associated with 318 assay endpoints across 103 tattoo ink chemicals (Table S8).

**7.2. Identification of KEs using CTD**

The Comparative Toxicogenomics Database (CTD) is a comprehensive public resource that links environmental chemicals (C), genes (G), phenotypes (P), and diseases (D) to advance understanding of their effects on health.[26] This resource facilitates the construction of CGPD-tetramers, which help identify confident associations between chemicals, phenotypes, and diseases, enabling their mapping to KEs within AOP-Wiki. Based on our previous work,[20–22] high confidence CGPD-tetramers associated with tattoo ink chemicals were retrieved, and subsequently leveraged to identify KEs within AOP-Wiki.

First, CTD May 2025 release was accessed and the CGPD-tetramers were constructed, wherein the following were considered: (i) chemical-gene and chemical-phenotype associations with literature evidence; (ii) chemical-disease and gene-disease associations with 'marker/mechanism' evidence; (iii) gene-phenotype associations with GO annotations based on only the experimental results (https://geneontology.org/docs/guide-go-evidence-codes/). This process resulted in a list of 59342 CGPD-tetramers comprising 63 tattoo ink chemicals, 2397 genes, 797 phenotypes and 502 diseases (Table S12). Next, the immediate neighboring GO terms for the CGPD-tetramer phenotype GO terms were generated using the GOSim package[36] in R programming language. These GO terms were then overlapped with the process identifiers of KEs in AOP-Wiki, and manually screened to identify 246 KEs linked to 176 phenotypes across 62 tattoo ink chemicals (Table S8). Additionally, the disease terms were manually mapped to identify 213 KEs associated with 406 diseases across 63 tattoo ink chemicals (Table S8).

##### 7.3. Identification of KEs using DEDuCT and NeurotoxKb

DEDuCT[27,28] (https://cb.imsc.res.in/deduct/) is one of the largest databases compiling curated information on endocrine disrupting chemicals (EDCs) and their corresponding endocrine-mediated endpoints from published literature. In this study, the endocrine-mediated endpoints corresponding to tattoo ink chemicals within DEDuCT were extracted and subsequently considered to identify relevant KEs within AOP-Wiki. The endocrine-mediated endpoints and KE titles in AOP-Wiki were manually inspected, to identify 142 KEs associated with 106 endocrine-mediated endpoints across 29 chemicals (Table S8).

NeurotoxKb[29] (https://cb.imsc.res.in/neurotoxkb/) is a manually curated resource focusing on mammalian neurotoxicity endpoints associated with environmental chemicals, compiled from published literature. In this study, the neurotoxic endpoints corresponding to tattoo ink chemicals within NeurotoxKb were extracted, and subsequently considered to identify relevant KEs within AOP-Wiki. The neurotoxic endpoints and KE titles in AOP-Wiki were manually inspected, to identify 34 KEs associated with 29 neurotoxic endpoints across 17 chemicals (Table S8).

##### 7.4. Identification of KEs using AOP-Wiki

AOP-Wiki catalogs information on prototypical stressors for each AOP, based on well-documented associations with these stressors (https://aopwiki.org/handbooks/5). In this study, the information on prototypical stressors associated with AOPs was extracted from the flat download file using an in-house Python script. Thereafter, 7 tattoo ink chemicals were identified as stressors for 18 AOPs. Subsequently, the 82 KEs within these AOPs were considered as associated with these tattoo chemicals (Table S8).

##### 7.5. Identification of KEs using REACH Dossier experimental information

The European chemicals agency (ECHA) provides a public database of chemicals that are registered in REACH (https://chem.echa.europa.eu/). In particular, various chemical information, including chemical properties, and experimental results from different toxicological studies have been curated into a dossier for each of the chemicals. In this study, the experimental toxicological information pertaining to tattoo chemicals were obtained from the publicly available chemical dossiers accessible at: https://chem.echa.europa.eu/ (last accessed on 3 July 2025) (Table S4), and subsequently leveraged to identify relevant KEs within AOP-Wiki.

First, reliable experimental information, pertaining to the different adverse effects associated with tattoo ink chemicals, were identified by relying on the studies marked as 'Key study', type of information as 'experimental study', and a reliance score denoted by Klimisch score[12] of 1 or 2. Next, mammalian specific genotoxic endpoints were identified based on the experimental design detailed in the dossiers. Thereafter, the different adverse effects and the genotoxic endpoints were manually inspected to identify 60 KEs associated with 84 adverse effects across 54 tattoo ink chemicals (Table S8).

Overall, 648 KEs were identified to be associated with 154 tattoo ink chemicals through the integration of heterogenous toxicogenomic and biological endpoints information from six exposome-relevant resources, namely, ToxCast, CTD, DEDuCT, NeurotoxKb, AOP-Wiki and REACH Dossiers (Table S8).

**8. Construction of stressor-AOP network**

Stressor-AOP network provides a broader perspective on impacts of stressors across diverse biological processes by linking the stressors to AOPs.[21,31] To better understand the perturbances caused by tattoo ink chemicals, a stressor-AOP network was constructed, which is a bipartite graph that linked tattoo ink chemicals to different AOPs within AOP-Wiki. In order to obtain high confidence associations between the chemicals and AOPs, only the curated list of 385 curated AOPs were relied upon (Table S5).

First, the tattoo chemicals were linked to any AOP if they shared at least one common KE. Thereafter, these links were characterized based on two criteria namely, the coverage score and the level of relevance. The coverage score of a stressor-AOP link is defined as the ratio of number of KEs within that AOP associated with the stressor to the total number of KEs within that AOP.[20,37] This score is a real valued number between 0 and 1, and is denoted as the edge weight of linkage between a stressor and an AOP in the constructed stressor-AOP network. The level of relevance is a qualitative score used to identify the relevance of stressor-AOP association within the stressor-AOP network,[21] and this score is denoted as an attribute of the edge in the constructed stressor-AOP network. Level of relevance is a five-level criterion defined as follows:

- *Level 1*: The stressor is associated with at least one KE within an AOP, where the KE is neither MIE nor AO within that AOP

- *Level 2*: The stressor is associated with at least one AO within an AOP, but not associated with any MIE within that AOP
- *Level 3*: The stressor is associated with at least one MIE within an AOP, but not associated with any AO within that AOP
- *Level 4*: The stressor is associated with at least one MIE and one AO within an AOP
- *Level 5*: The stressor is associated with at least one MIE and one AO within an AOP and there exists a directed path between the associated MIE and AO

Table S9 contains all the data on the stressor-AOP network constructed for tattoo ink chemicals, including the coverage score and level of relevance for each of the stressor-AOP links.

**References**

(1) Mohammed Taha, H.; Aalizadeh, R.; Alygizakis, N.; Antignac, J.-P.; Arp, H. P. H.; Bade, R.; Baker, N.; Belova, L.; Bijlsma, L.; Bolton, E. E.; Brack, W.; Celma, A.; Chen, W.-L.; Cheng, T.; Chirsir, P.; Čirka, Ľ.; D'Agostino, L. A.; Djoumbou Feunang, Y.; Dulio, V.; Fischer, S.; Gago-Ferrero, P.; Galani, A.; Geueke, B.; Głowacka, N.; Glüge, J.; Groh, K.; Grosse, S.; Haglund, P.; Hakkinen, P. J.; Hale, S. E.; Hernandez, F.; Janssen, E. M.-L.; Jonkers, T.; Kiefer, K.; Kirchner, M.; Koschorreck, J.; Krauss, M.; Krier, J.; Lamoree, M. H.; Letzel, M.; Letzel, T.; Li, Q.; Little, J.; Liu, Y.; Lunderberg, D. M.; Martin, J. W.; McEachran, A. D.; McLean, J. A.; Meier, C.; Meijer, J.; Menger, F.; Merino, C.; Muncke, J.; Muschket, M.; Neumann, M.; Neveu, V.; Ng, K.; Oberacher, H.; O'Brien, J.; Oswald, P.; Oswaldova, M.; Picache, J. A.; Postigo, C.; Ramirez, N.; Reemtsma, T.; Renaud, J.; Rostkowski, P.; Rüdel, H.; Salek, R. M.; Samanipour, S.; Scheringer, M.; Schliebner, I.; Schulz, W.; Schulze, T.; Sengl, M.; Shoemaker, B. A.; Sims, K.; Singer, H.; Singh, R. R.; Sumarah, M.; Thiessen, P. A.; Thomas, K. V.; Torres, S.; Trier, X.; van Wezel, A. P.; Vermeulen, R. C. H.; Vlaanderen, J. J.; von der

Ohe, P. C.; Wang, Z.; Williams, A. J.; Willighagen, E. L.; Wishart, D. S.; Zhang, J.; Thomaidis, N. S.; Hollender, J.; Slobodnik, J.; Schymanski, E. L. The NORMAN Suspect List Exchange (NORMAN-SLE): Facilitating European and Worldwide Collaboration on Suspect Screening in High Resolution Mass Spectrometry. *Environmental Sciences Europe* **2022**, *34* (1), 104. https://doi.org/10.1186/s12302-022-00680-6.

(2) Williams, A. J.; Grulke, C. M.; Edwards, J.; McEachran, A. D.; Mansouri, K.; Baker, N. C.; Patlewicz, G.; Shah, I.; Wambaugh, J. F.; Judson, R. S.; Richard, A. M. The CompTox Chemistry Dashboard: A Community Data Resource for Environmental Chemistry. *Journal of Cheminformatics* **2017**, *9* (1), 61. https://doi.org/10.1186/s13321-017-0247-6.

(3) Grulke, C. M.; Williams, A. J.; Thillanadarajah, I.; Richard, A. M. EPA's DSSTox Database: History of Development of a Curated Chemistry Resource Supporting Computational Toxicology Research. *Computational Toxicology* **2019**, *12*, 100096. https://doi.org/10.1016/j.comtox.2019.100096.

(4) Dionisio, K. L.; Phillips, K.; Price, P. S.; Grulke, C. M.; Williams, A.; Biryol, D.; Hong, T.; Isaacs, K. K. The Chemical and Products Database, a Resource for Exposure-Relevant Data on Chemicals in Consumer Products. *Scientific Data* **2018**, *5* (1), 180125. https://doi.org/10.1038/sdata.2018.125.

(5) Piccinini, P.; Contor, L.; Pakalin, S.; Raemaekers, T.; Senaldi, C. *Safety of Tattoos and Permanent Make-up – State of Play and Trends in Tattoo Practices*; EUR 27528; Publications Office of the European Union, JRC96808: Luxembourg, 2015. https://publications.jrc.ec.europa.eu/repository/handle/JRC96808.

(6) National Industrial Chemicals Notification and Assessment Scheme (NICNAS); Australian Government, Department of Health. *Characterisation of Tattoo Inks Used in*

*Australia*; 2018. https://www.industrialchemicals.gov.au/consumers-and-community/tattoo-and-permanent-make-pmu-inks.

(7) National Industrial Chemicals Notification and Assessment Scheme (NICNAS); Australian Government, Department of Health. *Investigation of the Composition and Use of Permanent Make-up (PMU) Inks in Australia*; 2017. https://www.industrialchemicals.gov.au/consumers-and-community/tattoo-and-permanent-make-pmu-inks.

(8) Hunger, K.; Schmidt, M. U. *Industrial Organic Pigments: Production, Crystal Structures, Properties, Applications*; Wiley-VCH Verlag, 2018.

(9) Hunger, K. *Industrial Dyes: Chemistry, Properties, Applications*; Wiley-VCH Verlag, 2002.

(10) Buxbaum, G.; Pfaff, G. *Industrial Inorganic Pigments*; Wiley-VCH Verlag, 2005.

(11) OECD. *Test No. 105: Water Solubility*; OECD Guidelines for the Testing of Chemicals, Section 1; OECD, 1995.

(21) Sahoo, A. K.; Chivukula, N.; Madgaonkar, S. R.; Ramesh, K.; Marigoudar, S. R.; Sharma, K. V.; Samal, A. Leveraging Integrative Toxicogenomic Approach towards

Development of Stressor-Centric Adverse Outcome Pathway Networks for Plastic Additives. *Archives of Toxicology* **2024**, *98* (10), 3299–3321. https://doi.org/10.1007/s00204-024-03825-z.

(27) Karthikeyan, B. S.; Ravichandran, J.; Mohanraj, K.; Vivek-Ananth, R. P.; Samal, A. A Curated Knowledgebase on Endocrine Disrupting Chemicals and Their Biological

Systems-Level Perturbations. *Science of The Total Environment* **2019**, *692*, 281–296. https://doi.org/10.1016/j.scitotenv.2019.07.225.

### Supplementary Figures

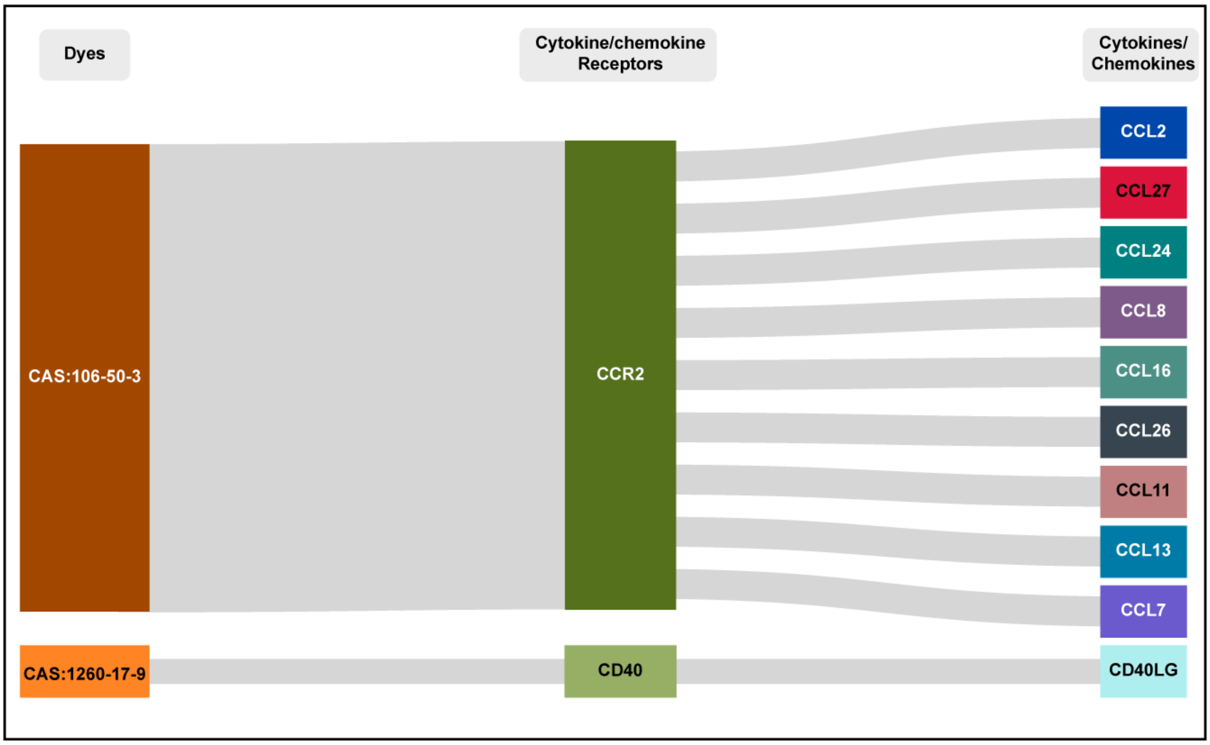


**Figure S1:** Visualization of the tattoo ink chemical - cytokine/chemokine receptors - cytokines/chemokines tripartite network, comprising two tattoo ink chemicals classified as ‘Dyes’.

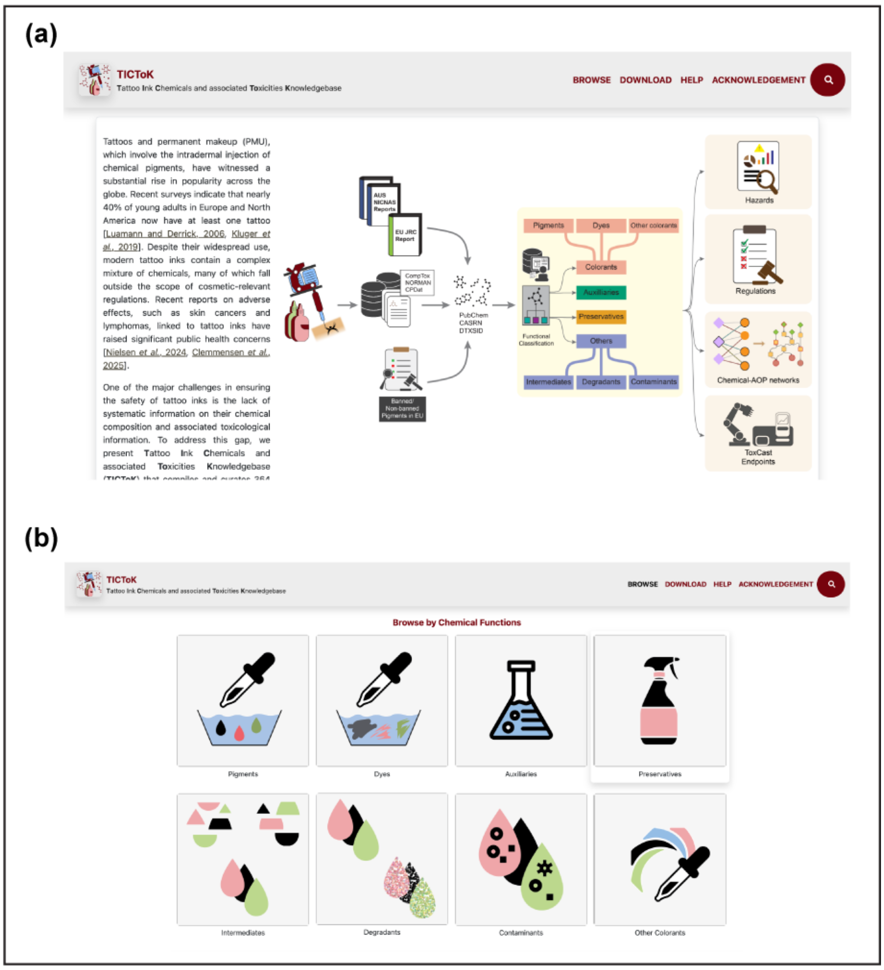


**Figure S2**: TICToK website. (a) The home page of TICToK web server with an easy navigation bar. (b) Browse page for chemicals based on their function in tattoo inks.

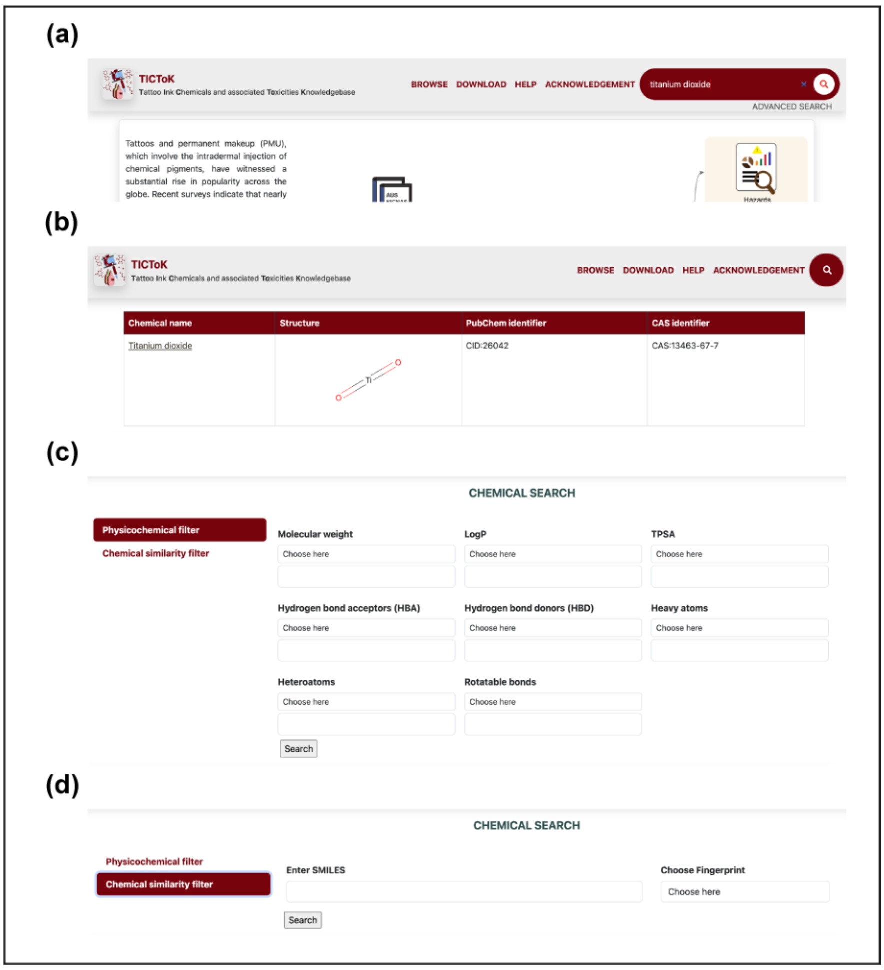


**Figure S3:** TICToK images of the 'SEARCH' sections. (a) The chemicals can be searched through their chemical identifiers from the top right corner of the page. (b) The search results in a tabulated page that contains different information pertaining to the search term. (c) ADVANCED SEARCH option based on physicochemical filtrations. (d) ADVANCED SEARCH option based on chemical similarity.

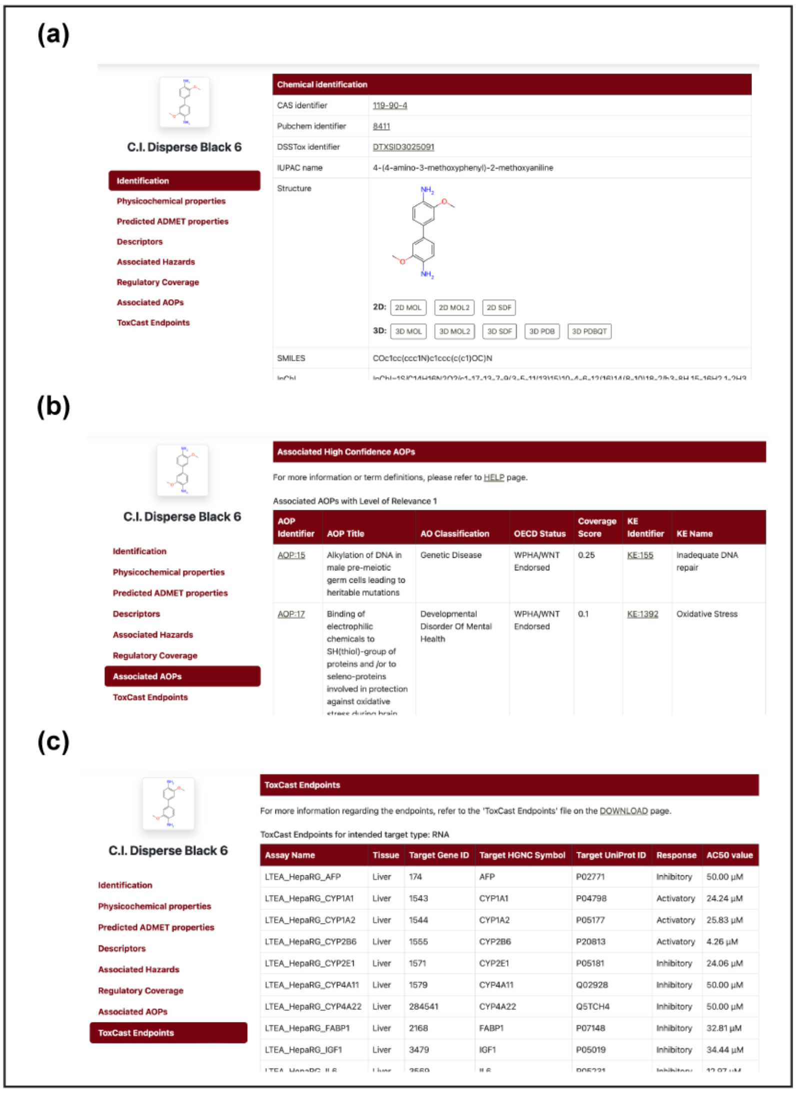


**Figure S4**: TICToK images of CHEMCIAL INFORMATION page. (a) Chemical identification page containing various chemical information and structure download files. (b) 'Associated AOPs' section containing information on the AOPs associated with the chemical. (c) 'ToxCast Endpoints' section contains the active toxicological endpoints associated with the chemical from ToxCast invitrodb v4.2.
